# Structure of the RZZ complex and molecular basis of Spindly-driven corona assembly at human kinetochores

**DOI:** 10.1101/2021.12.03.471119

**Authors:** Tobias Raisch, Giuseppe Ciossani, Ennio d’Amico, Verena Cmentowski, Sara Carmignani, Stefano Maffini, Felipe Merino, Sabine Wohlgemuth, Ingrid R. Vetter, Stefan Raunser, Andrea Musacchio

**Author notes:** These authors contributed equally (Tobias Raisch, Giuseppe Ciossani, Ennio d’Amico, Verena Cmentowski). European Institute of Oncology, Via Adamello 16, 20100 Milan, Italy (Giuseppe Ciossani). Max Planck Institute for Developmental Biology, Department of Protein Evolution, Max-Planck-Ring 5, 72076 Tübingen, Germany (Felipe Merino).

## Abstract

In metazoans, a ≍1 megadalton (MDa) super-complex comprising the Dynein-Dynactin adaptor Spindly and the ROD-Zwilch-ZW10 (RZZ) complex is the building block of a fibrous biopolymer, the kinetochore fibrous corona. The corona assembles on mitotic kinetochores to promote microtubule capture and spindle assembly checkpoint (SAC) signaling. We report here a high-resolution cryo-EM structure that captures the essential features of the RZZ complex, including a farnesyl binding site required for Spindly binding. Using a highly predictive *in vitro* assay, we demonstrate that the SAC kinase MPS1 is necessary and sufficient for corona assembly at supercritical concentrations of the RZZ-Spindly (RZZS) complex, and describe the molecular mechanism of phosphorylation-dependent filament nucleation. We identify several structural requirements for RZZS polymerization in rings and sheets. Finally, we identify determinants of kinetochore localization and corona assembly of Spindly. Our results describe a framework for the long-sought-for molecular basis of corona assembly on metazoan kinetochores.

## Introduction

Kinetochores are multi-subunit macromolecular assemblies that promote the bi-orientation and segregation of chromosomes during cell division (Musacchio and Desai, 2017; Navarro and Cheeseman, 2021). They are multi-layered structures built on specialized chromatin loci named centromeres. The kinetochore’s inner layer, named the constitutive centromere-associated network (CCAN) binds directly to the centromeric chromatin. The kinetochore’s outer layer, named the <u>K</u>nl1 complex, <u>M</u>is12 complex, <u>N</u>dc80 complex (KMN) network, generates a microtubule-binding interface. Additionally, the KMN network is a functional platform for the recruitment of several proteins that regulate the process of chromosome bi-orientation. Among the latter are proteins participating in the spindle assembly checkpoint (SAC), a cell cycle checkpoint that prevents mitotic exit before completion of bi-orientation (Lara-Gonzalez et al., 2021). SAC coordination is essential to prevent premature exit from mitosis or meiosis, which, by causing loss of chromosome cohesion in presence of unattached chromosomes, prevents successful chromosome partition to the daughter cells and is therefore essential for genome stability (Lara-Gonzalez et al., 2021).

In metazoans, including humans, checkpoint activity is coupled with the assembly of an outermost kinetochore layer named the kinetochore corona (Cooke et al., 1997; Hoffman et al., 2001; Jokelainen, 1967; Magidson et al., 2015; McEwen et al., 1993; Rieder and Alexander, 1990; Yao et al., 1997). The corona is a fibrous crescent-shaped structure that is only formed transiently in prometaphase cells, before the achievement of end-on attachment of chromosomes to microtubules (Kops and Gassmann, 2020). In checkpoint arrested cells, for instance in cells treated with agents, such as nocodazole, that promote microtubule depolymerization, the corona assumes a very characteristic expanded crescent shape that surrounds the core kinetochore and often even fuses with the corona nucleated by the sister kinetochore (Hoffman et al., 2001; Magidson et al., 2015; Pereira et al., 2018; Sacristan et al., 2018; Wynne and Funabiki, 2015).

In recent years, there has been substantial progress on the investigation of the mechanisms of assembly and disassembly of the kinetochore corona, and of its contributions to microtubule binding and SAC regulation (Kops and Gassmann, 2020). Proteins whose recruitment to kinetochore has been associated with assembly of the corona include the 3-subunit ROD-Zwilch-ZW10 complex (named after the *Drosophila melanogaster* genes *Rough Deal*, *Zwilch*, and *Zeste White 10*, and abbreviated as RZZ), the microtubule plus-end directed motor CENP-E, the microtubule-binding protein CENP-F, a tight “core complex” of the SAC proteins MAD1 and MAD2, and the microtubule minus-end directed motor Dynein, in complex with its processivity factor Dynactin (Dynein-Dynactin will be abbreviated as DD) and with a DD adaptor named Spindly (Karess, 2005; Kops and Gassmann, 2020).

Microtubule motors in the corona facilitate the process of chromosome alignment at the metaphase plate. After initial microtubule capture, these motors coordinate minus- and plus-end-directed transport of chromosomes that promotes their alignment at the metaphase plate before conversion of kinetochore attachment from lateral (i.e. to the microtubule lattice) to end-on (i.e. into the kinetochore interface). This conversion engages the core microtubule receptor of the kinetochore, the NDC80 complex, a sub-complex of the KMN network. A crucial aspect of the lateral to end-on conversion is that it coincides with a sudden activation of DD and with the disassembly of the kinetochore corona, in a process traditionally known as “shedding” (Auckland et al., 2020; Basto et al., 2004; Howell et al., 2001; Mische et al., 2008; Sivaram et al., 2009; Varma et al., 2008; Williams et al., 1996; Wojcik et al., 2001).

Corona shedding also coincides with silencing of SAC signaling at the particular kinetochore undergoing conversion to end-on attachment (Kuhn and Dumont, 2017; Kuhn and Dumont, 2019). The corona promotes the SAC by providing a docking site for the recruitment of the MAD1:MAD2 core complex, which is crucially required for checkpoint signaling (De Antoni et al., 2005; Faesen et al., 2017). SAC silencing is therefore caused by the removal, during corona shedding, of the MAD1:MAD2 core complex, which ultimately suppresses catalytic assembly of the checkpoint effector, the mitotic checkpoint complex (MCC) (Allan et al., 2020; Basto et al., 2000; Buffin et al., 2005; Caldas et al., 2015; Fava et al., 2011; Jackman et al., 2020; Kops et al., 2005; Matson and Stukenberg, 2014; Rodriguez-Rodriguez et al., 2018; Silio et al., 2015; Zhang et al., 2015).

The 812-kDa RZZ complex, whose subunits are shown schematically in Figure 1A, is a 2:2:2 hexamer (Civril et al., 2010; Mosalaganti et al., 2017; Scaerou et al., 2001). The RZZ is considered the corona’s building block (Mosalaganti et al., 2017; Pereira et al., 2018; Sacristan et al., 2018). While there is considerable interest in understanding how the RZZ promotes corona assembly, there is only limited structural insight into this process. An early structural analysis revealed the crystal structure of Zwilch and identified the 2209-residue ROD protein as a member of a family of proteins, which also includes clathrin, consisting of an N-terminal *β*-propeller followed by a long *α*-solenoid (Civril et al., 2010). Reconstitution of the RZZ and a single particle electron cryo microscopy (cryo-EM) structure, limited to an average resolution of approximately 10-12 Å, offered the first comprehensive view of the organization of the RZZ hexamer (Altenfeld et al., 2015; Mosalaganti et al., 2017). The reconstruction demonstrated that two highly elongated ROD protomers are arranged in an anti-parallel configuration and that a ZW10 dimer cements this organization in its central region, while Zwilch occupies a more peripheral position, between ZW10 and the ROD *β*-propeller (Mosalaganti et al., 2017). Also based on homologous proteins of known structure but very limited sequence homology, structural models were built for ROD and ZW10 to fit the EM reconstruction (Mosalaganti et al., 2017). This previous work, however, failed to provide a detailed molecular description of the RZZ subunits and of their interactions. Here, we fill this gap by reporting a high-resolution structure of the RZZ complex, obtained by single-particle cryo-EM that finally reveals all its detailed structural features.

**Figure 1.**
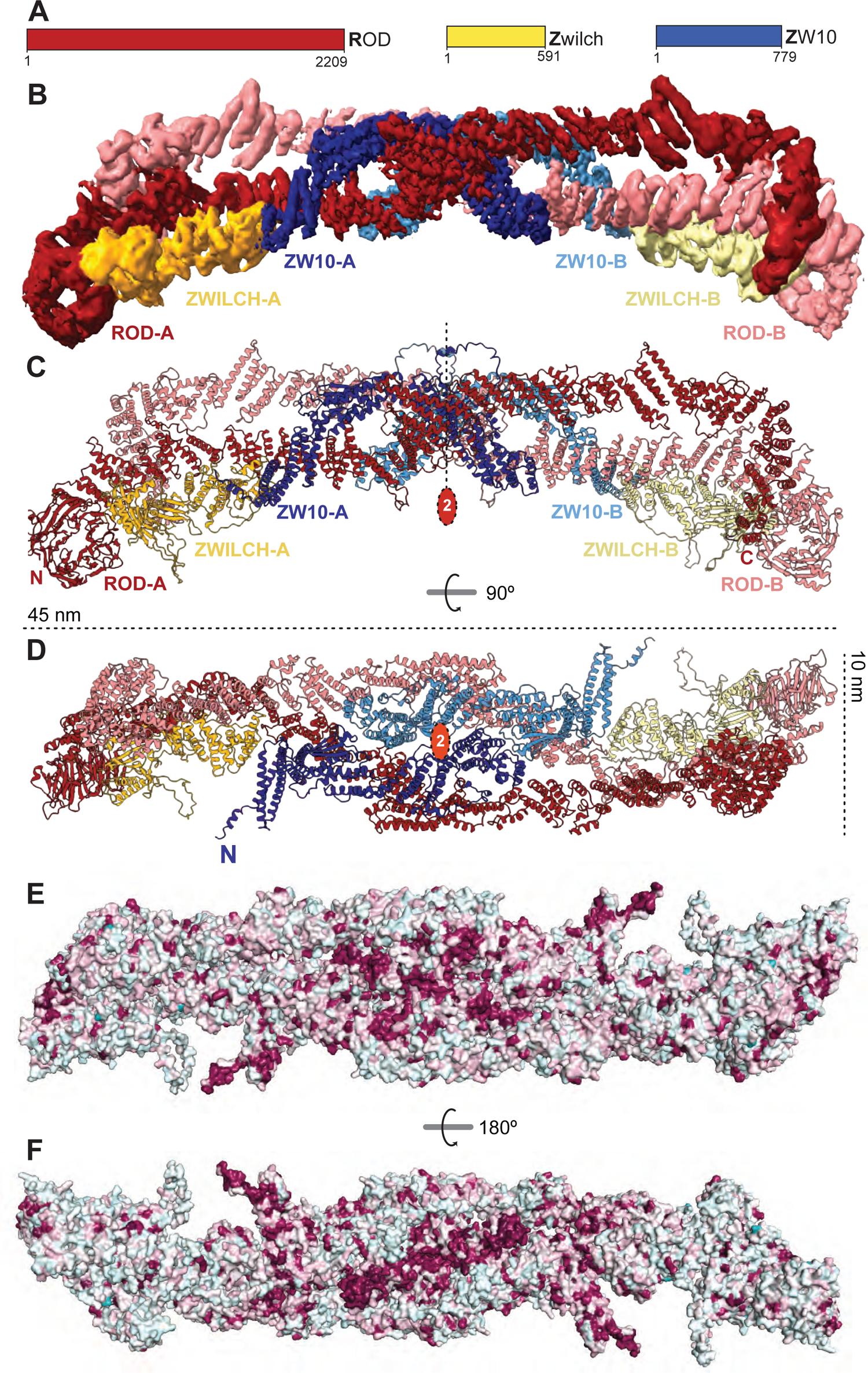
Structural organization of the RZZ complex. (**A**) Schematic representation of human RZZ subunits. (**B**) 3D reconstruction of the RZZ complex with densities corresponding to ROD-A, Zwilch-A, and ZW10-A colored in firebrick, yellow-orange, and deepblue, respectively. ROD-B, Zwilch-B, and ZW10-B are displayed in equivalent lighter colors as indicated. (**C**) Cartoon model of the RZZ complex with coloring scheme like in panel A. The position of the internal 2-fold axis of the 2:2:2 hexamer is shown. The N- and C-termini of ROD are indicated. Panels B-E and all other panels displaying molecular features were generated with PyMol (The PyMOL Molecular Graphics System, Version 1.2r3pre, Schrödinger, LLC.). (**D**) 90-degree-rotated view of the complex with linear dimensions. (**E**) The conservation of residues in an alignment of ROD, Zwilch, and ZW10 is displayed on the surface of the complex (dark, highly conserved; light, poorly conserved). For all subunits, conservation was calculated from an alignment of sequences from *C. elegans* (Ce), *D. melanogaster* (Dm), *X. tropicalis* (Xt), *D. rerio* (Dr), *Bos taurus* (Bt), *Mus musculus* (Mm), and *Homo sapiens* (Hs). (**F**) 180-degree-rotated view of the complex. The highest degree of conservation is observed in ZW10. See also alignments in Figure 1 – Supplement 1.

Spindly is a member of a large family of DD adaptors (Reck-Peterson et al., 2018) shown to activate dynein motility *in vitro* (Cianfrocco et al., 2015; Gama et al., 2017; Hoogenraad and Akhmanova, 2016; McKenney et al., 2014; Pereira et al., 2018; Sacristan et al., 2018; Schlager et al., 2014). How Spindly coordinates its interaction with RZZ with activation of DD motility and processivity remains poorly understood. Spindly binds directly to the RZZ complex through its C-terminal region (forming the complex abbreviated as RZZS), and engaging an RZZ module comprising the ROD *β*-propeller and Zwilch (Gama et al., 2017; Henen et al., 2021; Mosalaganti et al., 2017; Pereira et al., 2018; Sacristan et al., 2018). Furthermore, in humans and likely most other metazoans, Spindly is post-translationally modified on Cys602 with farnesyl, an isoprenoid lipid. This modification, which is required for the interaction of Spindly with RZZ, may engage a dedicated binding site on the ROD *β*-propeller (Gama et al., 2017; Holland et al., 2015; Mosalaganti et al., 2017; Moudgil et al., 2015).

Besides interacting with RZZ, Spindly is also required for kinetochore recruitment of DD (Barisic et al., 2010; Chan et al., 2009; Cheerambathur et al., 2013; Gama et al., 2017; Gassmann et al., 2008; Gassmann et al., 2010; Griffis et al., 2007; Raaijmakers et al., 2013; Starr et al., 1998; Yamamoto et al., 2008). The determinants of RZZ binding and DD recruitment by Spindly are separable. A region of Spindly, the Spindly motif, shown to be a conserved feature of adaptors, can be mutated to abrogate kinetochore recruitment of DD (Gama et al., 2017; Gassmann et al., 2010; Pereira et al., 2018; Sacristan et al., 2018). The mutation is compatible with corona expansion and chromosome bi-orientation, but preventing DD recruitment leads to a permanent SAC arrest caused by the inability to disassemble (strip) the corona and silence the SAC (Gama et al., 2017; Gassmann et al., 2010; Pereira et al., 2018; Sacristan et al., 2018).

Initial studies in humans and *C. elegans* identified conditions *in vitro* and in living cells for RZZ assembly into filamentous structural mimics of the corona, pointing to the RZZ as a candidate building block of the corona (Henen et al., 2021; Pereira et al., 2018; Sacristan et al., 2018). These recent studies, however, also brought to light different minimal requirements for filament assembly (Pereira et al., 2018; Sacristan et al., 2018), with species-specific differences and a persisting question on whether Spindly is necessary for filament assembly and acts as gatekeeper in this process. Corona assembly is limited to kinetochores and is sensitive to the cellular concentration of RZZ (Pereira et al., 2018), suggesting it requires a critical concentration that is exclusively reached upon RZZ recruitment to kinetochores in early prometaphase. Kinetochore recruitment, however, is not sufficient for corona assembly in human cells, because the depletion of Spindly or the inhibition of the SAC kinase MPS1 prevent expansion of the corona without preventing kinetochore recruitment of the RZZ(Pereira et al., 2018; Rodriguez-Rodriguez et al., 2018). Here, we have recapitulated with purified components *in vitro* the requirement for human ROD phosphorylation by MPS1 and Spindly binding for corona assembly. This assay allowed us to identify various additional requirements for nucleation of filaments by the RZZS complex, and to acquire structural information on the mechanism of filament assembly that was related to the high-resolution structure of the RZZ. We present a model for corona assembly whose implications were extensively corroborated with experiments in mitotic cells. Collectively, our results greatly advance our understanding of a fundamental aspect of kinetochore structure and function.

## Results

### Reconstitution and structural analysis of the RZZ complex

Using reconstituted human RZZ (Altenfeld et al., 2015; Mosalaganti et al., 2017), we previously reported a single particle cryo-EM reconstruction of the RZZ at an overall resolution of ∼10-12 Å (1 Å = 0.1 nm). As only the structure of Zwilch (Civril et al., 2010) (PDB ID 3IF8) had been experimentally determined, we had tried to account for the observed density by building *ad hoc* homology models of ROD or ZW10 and fitting them in the 3D reconstruction (Mosalaganti et al., 2017). Due to the very low resolution of the reconstructions, however, the resulting models were merely tentative.

To improve the resolution of the RZZ structure, we used an mCherry-tagged RZZ construct (^mCh^RZZ) that was better expressed than the previously used poly-histidine-tagged RZZ (see Methods). In addition, purified ^mCh^RZZ proved to be more stable than the previous construct, allowing us to determine the structure of the complex by cryo-EM at a resolution of 3.9 Å (Figure 1B). The new reconstruction allowed us to build an essentially complete atomic model of ZW10 and the central region of ROD. In the periphery of RZZ, where the resolution of the reconstruction was lower than in the center and therefore did not allow unequivocal model building, we resorted to high-confidence AlphaFold2 (AF2; (Jumper et al., 2021; Tunyasuvunakool et al., 2021)) predictions, and used flexible fitting with minimal interventions, to fit them in the density (Figure 1 –Supplement 1).

While related to our previous low-resolution model in its general outline, the new model provides a detailed description of all crucial molecular features of the RZZ complex. The RZZ complex is a 2:2:2 hexamer with C2 symmetry. The 2-fold-related ROD chains (A and B) run in an anti-parallel configuration that sets the ∼45-nm long and ∼10 nm wide dimensions of the RZZ (Figure 1C-D). After an N-terminal 7-bladed *β*-propeller, the ROD chain transitions, near residue 395, into a short helical hairpin that begins an uninterrupted but irregular *α*-solenoid that extends until the C-terminus.

Several proteins share with ROD a succession of an N-terminal *β*-propeller followed by a C-terminal *α*-solenoid. These include Clathrin, COP1, the Nup155 and Nup145 nucleoporins, Sec31, Sec39, and even the APC1 subunit of the APC/C (Alfieri et al., 2017; Brohawn et al., 2008; Fath et al., 2007; Fotin et al., 2004; Lee and Goldberg, 2010; Stagg et al., 2006; Stagg et al., 2007; ter Haar et al., 1998; Watson et al., 2019). In comparison to the 1675-residue Clathrin heavy chain or the 1944-residue APC1, the 2209-residue ROD *α*-solenoid is significantly more elongated and straighter (Figure 2 – Supplement 1A). Packing of successive helical hairpins against each other with a slight right- or a left-handed rotation is a typical pattern of regular *α*-solenoids. This pattern is also observed for blocks of successive helical hairpins in ROD, but there are points where the hairpins rather pack almost at a right angle, deflecting the polypeptide chain (Figure 2 – Supplement 1A). There are at least three points where the ROD chain bends sharply, around residues 856 and 1060 in the central region, and around residue 1905 in the C-terminal region. The latter kink generates a characteristic C-terminal “hook” that is perpendicular to the opposite ROD chain (Figure 2A and Figure 2 – Supplement 1A). This is a prominent interaction interface between ROD-A and ROD-B, as residues 1790-2125, which encompass part of the hook domain, form a cradle that interacts with residues 505-690 of the opposing ROD protomer, including residues 655-680, situated in an inter-helical loop (Figure 2B).

**Figure 2.**
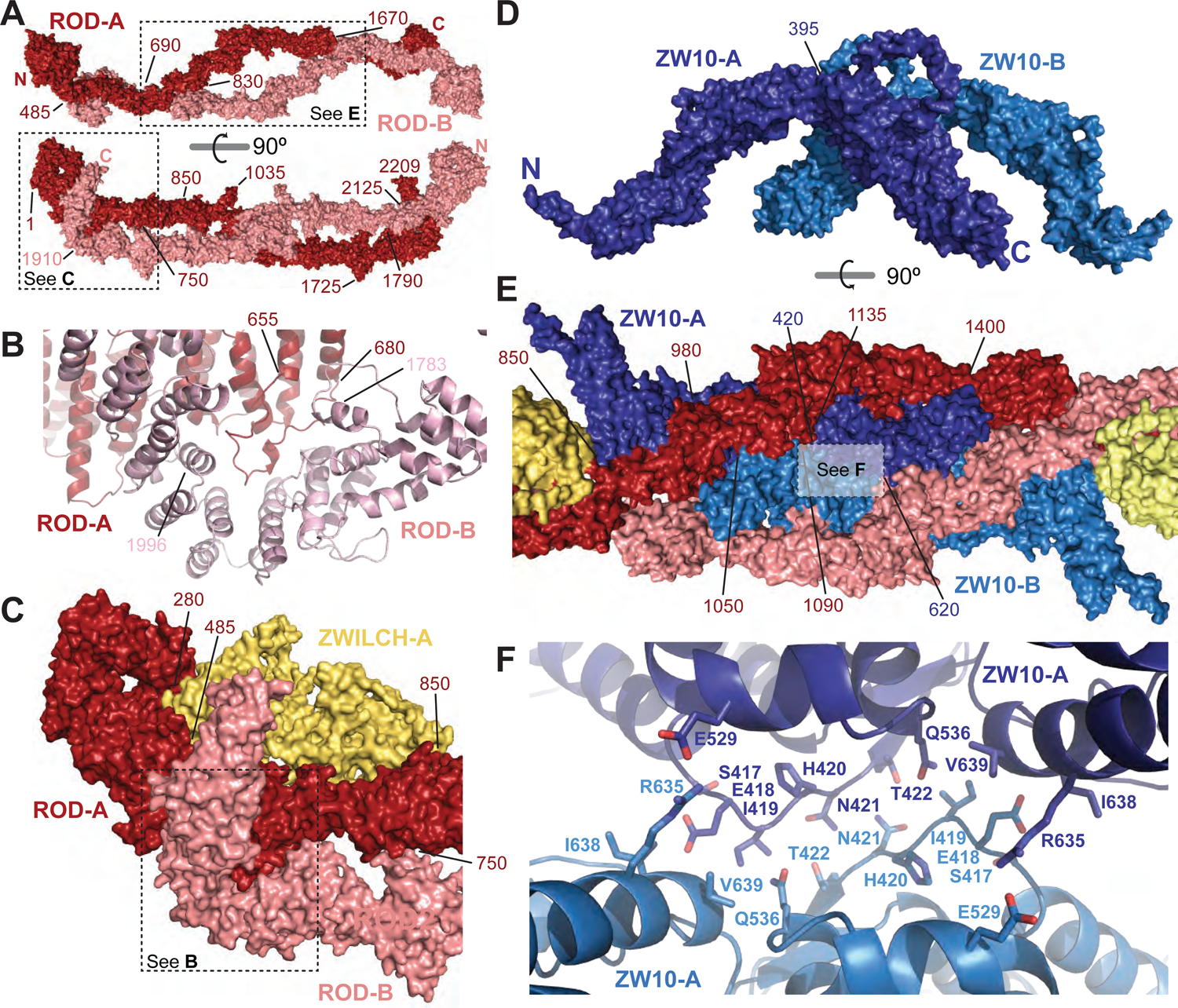
Homo- and heterotypic intermolecular interactions in RZZ. (**A**) The ROD-A and ROD-B protomers are related by 2-fold symmetry and interact through two main regions positioned between residues 485-690 and 1790-2125. (**B**) A particular prominent interaction of ROD-A and ROD-B consists in the insertion of the 655-680 loop of one protomer into a cradle formed by the bending C-terminal region of the second protomer. (**C**) Zwilch interacts very prominently with only one of the two ROD protomers, sandwiched between the N-terminal *β*-propeller and residue 850. A few interactions also link Zwilch to the C-terminal region of the second protomer. (**D**) A rotated view shows that ZW10 forms a highly bent, U-shaped complex with a relatively small inter-protomer interface. (**E**) The 2-fold axis of the complex (shown in Figure 1B) crosses the interface between ZW10-A and ZW10-B. (**F**) Molecular details of the interface between ZW10-A and ZW10-B with side chains of residues involved.

Zwilch abuts the ROD *β*-propeller nd forms a direct, extensive interface with only one of the ROD protomers, with only a small contact with the hook domain of the second ROD molecule (Figure 2C). Contacts of Zwilch with ROD terminate around residue 850 of ROD, where Zwilch also contacts ZW10 in a small 3-way interface. As expected, the structure of Zwilch in the RZZ complex is closely related to the crystal structure of Zwilch obtained in isolation (Civril et al., 2010), but adopts a more open conformation due to a reciprocal rotation of Zwilch’s two domains, presumably elicited by contacts within the complex (Figure 2 – Supplement 1B).

Finally, ZW10 adopts a highly curved, U-shaped conformation, a major determinant of which is the sharp bending around residue 395, situated between the N- and C-terminal domains (Figure 2D and Figure 2 – Supplement 1C). ZW10-related domains are found in Dsl1 and Tip20, subunits of vesicle tethering complexes in *S. cerevisiae*. Like ZW10, they both consist of two roughly equally sized helical domains (Tripathi et al., 2009), but are characterized by different inter-domain angles (Figure 2 – Supplement 1). Indeed, the isolated ZW10 shows high flexibility between its two domains (Mosalaganti et al., 2017).

In the RZZ, the interface between the A and B protomers of ZW10, which intersects the 2-fold symmetry axis of the RZZ complex, is relatively small (Figure 2D-F). Accordingly, AF2 does not predict any solitary ZW10 dimer in a conformation seen in the RZZ complex (unpublished results). ZW10 A and B, however, are stably set inside an “eye” between the two ROD chains (compare panels A and E in Figure 2), with which they form a very extensive interaction interface. Specifically, residues 850-980, 1050-1090, and 1135-1400 of ROD-A interact with ZW10-A, ZW10-B, and ZW10-A, respectively. The N-terminal regions of ZW10 are prominent features that emerge almost perpendicularly from the RZZ’s long axis. Together with the C-terminal region of ZW10, they are among the best conserved sequence features of the RZZ complex (Figure 1E-F; alignments are provided in Figure 1 – Supplement 2).

### ROD’s farnesyl binding site

The interaction of Spindly with RZZ is direct and requires isoprenylation of Spindly with a farnesyl group at Cys602 (Mosalaganti et al., 2017) (Figure 3A). Using various farnesyl moieties modified with photoactivatable cross-linkers and enzymatically incorporated in Spindly, we previously mapped a farnesyl binding site near Leu120 of ROD (Mosalaganti et al., 2017). This residue is located in proximity of a prominent feature of the ROD *β*-propeller, the insertion of an *α*-helical hairpin (residue 168-190) between strands *β*3C and *β*3D (Figure 3B and Figure 1 – Supplement 2). The hairpin abuts against blade 2 of the propeller, partly bending it and increasing its separation from blade 3, and generating a deep, roughly cylindrical cavity between the two blades (Figure 3C). Remarkably, Leu120 lines the entry point of the cavity. AF2 predicts the C-terminal region of Spindly to interact with this region of the ROD *β*-propeller, and modelling the farnesyl group on Cys602 shows that the cavity is ideally dimensioned to receive the farnesyl group (Figure 3D-G).

**Figure 3.**
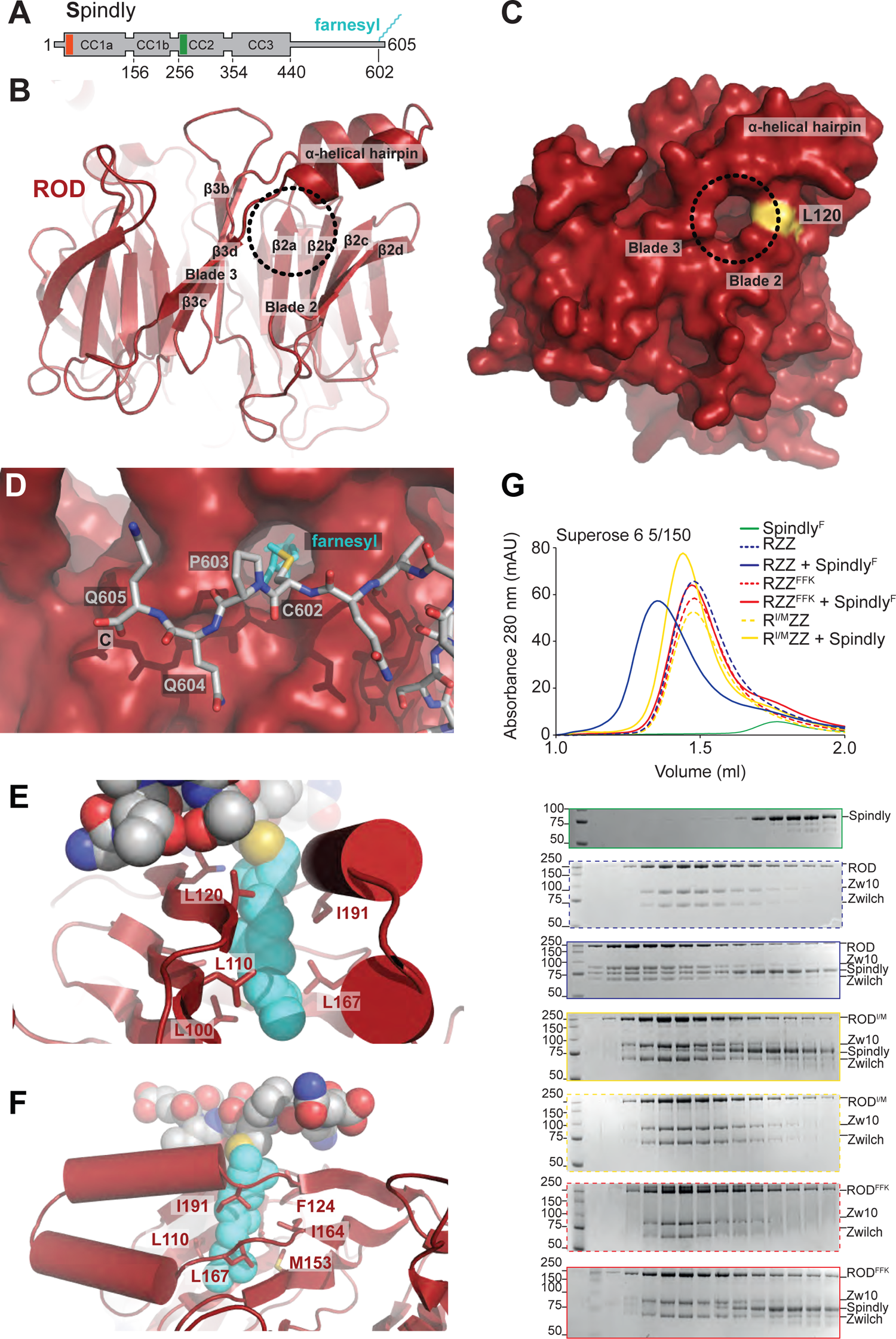
A farnesyl-binding pocket in the ROD propeller. (**A**) Schematic representation of HsSpindly. The position of a CC1 box (red) and of the Spindly box (green) required for binding Dynein:Dynactin are shown, together with the position of relevant predicted coiled-coil regions. (**B**) Cartoon representation of the 7-bladed ROD *β*-propeller (indicated as blades 1-7). The positions of blade 2, blade 3, and an *α*-helical hairpin representing an insertion between strands *β*C3 and *β*D3 are indicated. The four strands of each blade is indicated as A-D, with A and D being the innermost and outermost strands. The circle represents the entry point of the farnesyl binding pocket. (**C**) Surface representation of the ROD *β*-propeller. The circle is in the same position shown in panel A. The position of Leu120 is shown in yellow. This residue was was targeted by photoactivatable crosslinker groups introduced in the farnesyl group attached to Cys602 of Spindly (Mosalaganti et al., 2017). (**D**) The position of a modelled peptide corresponding to the C-terminal region of Spindly as predicted by AlphaFold2 (Jumper et al., 2021; Tunyasuvunakool et al., 2021) with a farnesyl moiety modelled on Cys602. (**E-F**) A farnesyl moiety, shown in cyan spheres together with a few C-terminal residues of a modelled Spindly peptide, fitted snugly into a pocket lined exclusively by the side chains of several hydrophobic residues. (**G**) Size-exclusion chromatography profiles and corresponding SDS-PAGE of the indicated samples. Two mutant RZZ complexes containing mutations in ROD L110F-L119F-L120K or I191M are indicated respectively as FFK and I/M. Note that in the bottom SDS-PAGE the first and second lane were deliberately inverted and contain the first eluted fraction and the molecular weight marker. For all other shown SDS-PAGE gels, the marker precedes the first fraction.

The entire cavity is lined with hydrophobic residues, including Leu100, Leu108, Leu110, Leu119, Leu120, Phe124, Met153, Ile164, Leu167, Leu169, and Ile191, in addition to two polar residues, Asn122 and Ser193 (Figure 3E-F). To test the role of this pocket in the binding of farnesylated Spindly (Spindly^F^), we tried to occlude it by generating two mutant RZZ complexes in which hydrophobic residues lining the farnesyl-binding pocket were replaced with bulkier ones (as described in the legend of Figure 3G). Confirming our hypothesis, the resulting mutants were stable but apparently unable to interact with Spindly^F^ in a size-exclusion chromatography experiment, contrarily to wild type RZZ, with which Spindly^F^ formed a stoichiometric complex (Figure 3G).

The *C. elegans* and *D. melanogaster* Spindly sequences have no C-terminal cysteine for isoprenylation (Holland et al., 2015). Analysis of ROD sequences in these organisms demonstrates differences predicted to ablate the hydrophobic farnesyl-binding cavity observed in human ROD. Specifically, in both species, the first of the two *α*-helices in the *α*-helical hairpin insertion that contributes to the architecture of the farnesyl-binding cavity is shorter by 3-residues (Figure 1 – Supplement 2). AF2 predicts that this causes a rotation of the second *α*-helix, positioning the side chain of Met184 (CeROD) precisely in the center of the cavity, obstructing it and making it inviable for farnesyl binding (Figure 3 – Supplement 1).

### MPS1 nucleates RZZS fibers

The requirements for corona expansion remain incompletely characterized. For instance, deletion of the ROD-1 *β*-propeller (residues 1-372) promotes ectopic filament formation in *C. elegans* embryos, while the equivalent deletion (residues 1-375) prevents expansion in human cells (Gama et al., 2017; Pereira et al., 2018). *In vitro*, a complex of CeZW10 (CZW-1) and CeROD with a deleted N-terminal propeller assembles a polymeric filamentous structure, whereas a full-length ternary complex is unable to polymerize (Pereira et al., 2018). Conversely, *in vitro* filament assembly of human RZZ, elicited by mild heating, requires Spindly^F^ (Sacristan et al., 2018). Furthermore, an N-terminal deletion removing 65 residues of Spindly promotes ectopic filament assembly in human cells even when Spindly is not farnesylated, an otherwise necessary condition for RZZ:Spindly binding and corona assembly (Sacristan et al., 2018).

To shed light on the corona assembly mechanism, we set out to dissect it *in vitro* with purified components. Using a spinning disk confocal microscope, we monitored polymerization of fluorescent mCherry-tagged RZZ or RZZS^F^ under various conditions after their purification to homogeneity and thorough dephosphorylation. At 20°C, neither ^mCh^RZZ nor ^mCh^RZZS^F^ (at a concentration of 4 µM) formed visible polymers (unpublished results and see below). Conversely, and in agreement with our previous observations (Sacristan et al., 2018), incubation of ^mCh^RZZS^F^ for 1 hour at 30°C promoted formation of copious fibers (Figure 4A). No fibers were observed with ^mCh^RZZ or ^mCh^Spindly^F^ (Figure 4A), indicating that Spindly^F^ is required for fiber formation.

**Figure 4.**
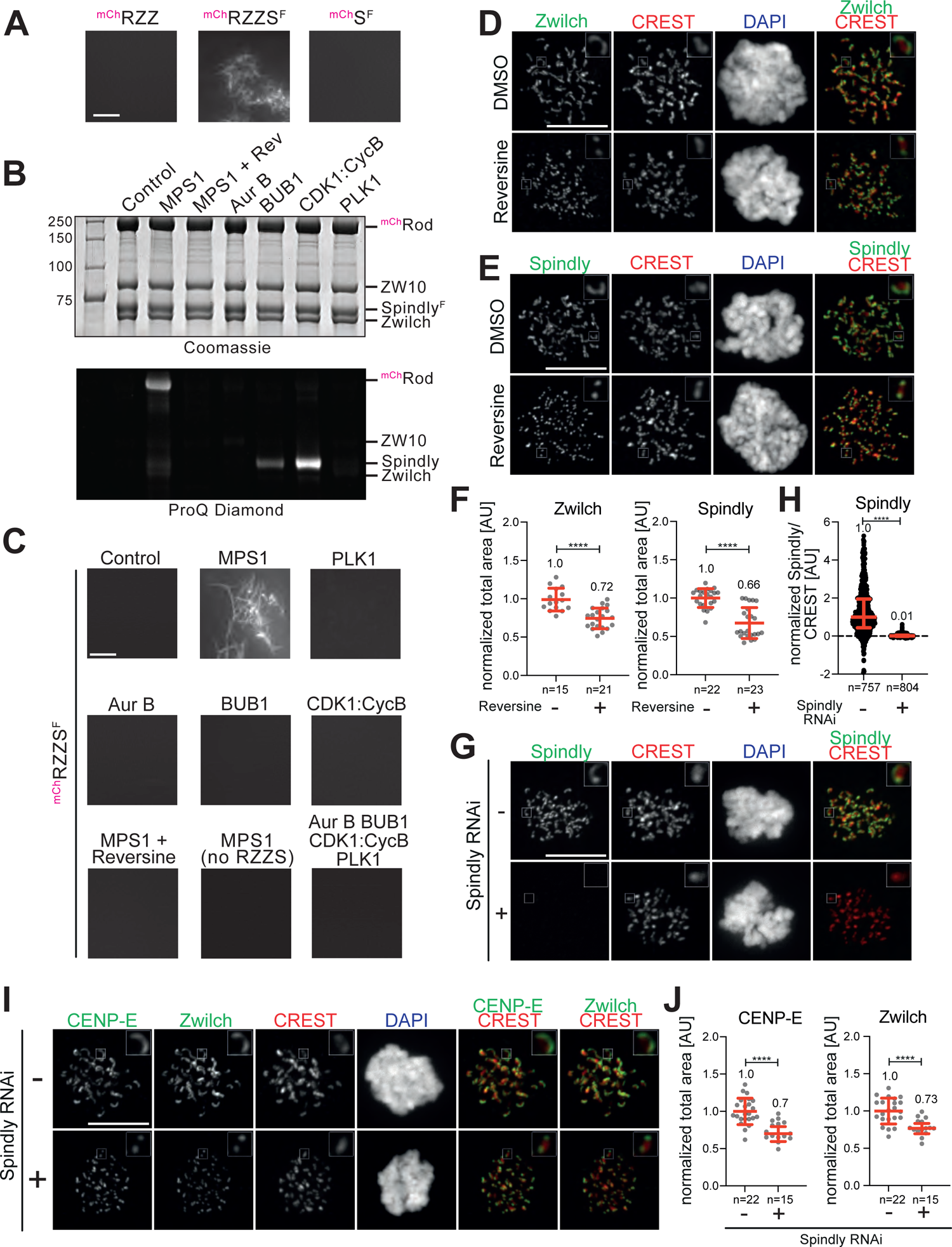
MPS1 and Spindly promote corona assembly. **(A)** Confocal fluorescence microscopy-based filamentation assay at 561 nm shows ^mCh^RZZS^F^ (4 µM), but not ^mCh^RZZ (4 µM) or ^mCh^S (4 µM) as controls, forms filaments at 30°C. Scale bar: 5 µm. (B) Coomassie and ProQ Diamond-stained gels on ^mCh^RZZS^F^ (4 µM) treated with the indicated kinases (1 µM) for 15 hours at 20°C. Rev = reversine, used at 10 µM. These were precisely the samples studied in experiments in panel C. (**C**) A filamentation assay as in **A** demonstrates sufficiency of MPS1 phosphorylation for filamentation. (**D**) Levels of Zwilch at kinetochores of HeLa cells that had been previously synchronized in G2 phase with 9 μM RO3306 for 16 h and then released into mitosis. Subsequently, cells were immediately treated with 500 nM Reversine, 3.3 µM nocodazole, and 10 µM MG132 for 1 hour and imaged while in mitosis. CREST serum was used to visualize kinetochores and DAPI to stain DNA. Scale bar: 10 µm. (**E**) Cells treated as for panel **D** were treated for visualization of Spindly. (**F**) Scatter dot plots representing normalized total area of the Zwilch and Spindly signals, normalized to the reversine-untreated control, in the indicated number of cells from the experiment shown in panels C-D. Red lines indicate mean and standard deviation. (**G**) Representative images showing the effects of a knockdown of the endogenous Spindly in HeLa cells. RNAi treatment was performed for 48 h with 50 nM siRNA. Before fixation, cells were synchronized in G2 phase with 9 μM RO3306 for 16 h and then released into mitosis. Subsequently, cells were immediately treated with 3.3 μM nocodazole for an additional hour. CREST serum was used to visualize kinetochores and DAPI to stain DNA. Scale bar: 10 µm (**H**) Scatter dot plots representing normalized intensity ratios of Spindly over CREST for individual kinetochores of cells from the experiment shown in panel **G**. Red lines indicate median with interquartile range. (**I**) Levels of CENP-E and Zwilch were assessed in control cells and in cells treated as in panel **G** to knockdown Spindly. (**J**) Scatter dot plots representing normalized total area of the CENP-E and Zwilch signals, normalized to the RNAi negative control, in the indicated number of cells from the experiment shown in panel I. Red lines indicate mean and standard deviation.

Previous studies identified a role of the MPS1 kinase, a central SAC component, in corona expansion in human cells (Rodriguez-Rodriguez et al., 2018; Sacristan et al., 2018). Regulatory effects of phosphorylation of corona components by additional mitotic kinases have also been reported (Allan et al., 2020; Barbosa et al., 2020; Pereira et al., 2018; Rodriguez-Rodriguez et al., 2018; Sacristan et al., 2018). We therefore asked if mitotic kinases influenced corona assembly in our assay *in vitro*. When subjected to sub-stoichiometric concentrations (typically 1/4 kinase/substrate ratio) of various mitotic kinases *in vitro* at 20°C in presence of ATP, ^mCh^RZZS^F^ sample was phosphorylated by MPS1 on ROD, and by CDK1:Cyclin-B and to a minor extent by BUB1 on Spindly^F^ (as visualized by staining with the Pro-Q^TM^ Diamond Gel Staining Reagent; Figure 4B).

In line with a role of MPS1 phosphorylation of ROD in corona assembly (Rodriguez-Rodriguez et al., 2018; Sacristan et al., 2018), addition of MPS1 and ATP promoted spontaneous assembly of fibers at 20°C, an effect that was eliminated in presence of reversine, an MPS1 inhibitor (Santaguida et al., 2010). Conversely, PLK1, Aurora B, BUB1, CDK1:Cyclin-B did not promote fiber assembly, even when combined (Figure 4C). Subjecting ^mCh^RZZ or Spindly^F^ to MPS1 phosphorylation before mixing them to form a complex showed that only ROD needs to be phosphorylated for fibers to assemble (Figure 4 – Supplement 1A-B). Spindly required farnesylation for filamentation because a C602A mutant failed to filament (Figure 4 – Supplement 1C).

To corroborate these results in an *in vivo* setting, we released HeLa cells from a G2-phase arrest into mitosis in presence of nocodazole and reversine to depolymerize microtubules and inhibit MPS1, respectively. MPS1 inhibition by reversine was confirmed by severe reduction of BUB1 kinetochore levels (Figure 4 – Supplement 1D-E), as reported previously (Santaguida et al., 2010). Both Zwilch and Spindly decorated kinetochores when MPS1 was inhibited, albeit at a slightly reduced level in comparison with the control condition in absence of MPS1 inhibitors (Figure 4D-E, quantified in panel F). However, corona expansion had been clearly completely inhibited in presence of reversine, because Zwilch and Spindly showed the same dotted appearance of inner kinetochore markers instead of the crescent-like appearance observed in control cells (Figure 4D-E).

Next, we asked if Spindly is necessary for corona expansion as predicted by our results *in vitro*. After depleting Spindly by RNAi (Figure 4G-H), we detected CENP-E and Zwilch at reduced but still highly significant levels on kinetochores of nocodazole-arrested mitotic cells. As for MPS1 inhibition, however, also in this case we observed an essentially complete failure to expand the corona (Figure 4I-J). Thus, neither Spindly nor MPS1 phosphorylation are strictly required for RZZ recruitment to the kinetochore. However, both are indispensable for expanding the corona.

Thus, the *in vitro* corona assembly assay we describe is an excellent predictor of cellular events at the corona, and leads to conclude that the presence of RZZ, Spindly, and active MPS1 are necessary to promote assembly of the corona at the kinetochore. Because corona assembly is limited to kinetochores, however, there must be additional signals to enrich the corona constituents to these subcellular compartments. When added to mitotic cells with an already formed corona, the selective cyclin-dependent kinase 1 (CDK1) inhibitor RO3306 (Vassilev et al., 2006) promoted corona detachment from the kinetochore, but no corona disassembly, as described previously (Pereira et al., 2018; Sacristan et al., 2018) (Figure 4 –Supplement 1F-G). Conversely, the selective Aurora B kinase inhibitor Hesperadin (Hauf et al., 2003) erased the recruitment of both RZZ (and therefore Spindly) and CENP-E to the kinetochore (Figure 4 – Supplement 1H). Thus, both CDK1 and Aurora B are essential for directing or retaining the corona to the kinetochore, but our results *in vitro* strongly suggest that they do so on the kinetochore side of the binding interface for the corona.

### The role of MPS1 phosphorylation

Thr13 and Ser15 of ROD were previously identified as MPS1 substrates required for corona assembly (Rodriguez-Rodriguez et al., 2018). In agreement with these results, we found that a ^mCh^RZZS^F^ mutant complex where ROD^Thr13^ and ROD^Ser15^ were mutated to alanine (T13A/S15A) did not assemble fibers (Figure 5A). The phosphomimetic mutant T13E/S15E, on the other hand, allowed ^mCh^RZZS^F^ to form fibers at 20°C and in presence of reversine to abrogate MPS1 activity (Figure 5A and Figure 5 – Supplement 1A-B). The T13E/S15E mutant, however, failed to form fibers when Spindly^F^ was omitted. This confirms that Spindly^F^ is absolutely necessary for fiber formation even after bypassing MPS1 activity. Collectively, these results confirm that our fiber formation assay *in vitro* recapitulates crucial aspects of corona assembly.

**Figure 5.**
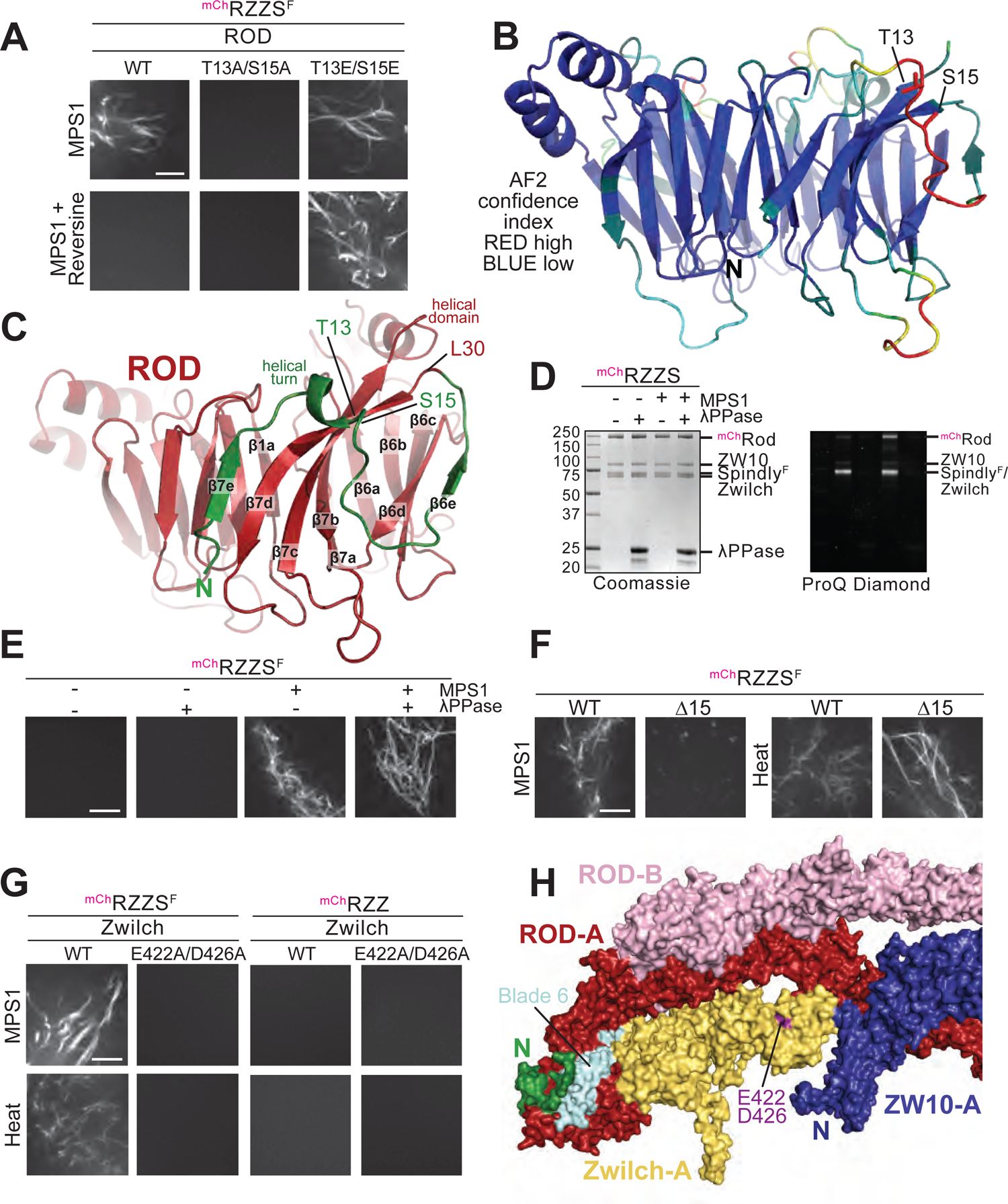
An autoinhibited state of the RZZS complex. (**A**) MPS1-induced filamentation experiments demonstrate that the phosphomimetic T13E/S15E ROD mutant bypasses the filamentation blockade induced by the MPS1 inhibitor reversine. The T13A/S15A mutant prevents MPS1-induced filamentation altogether. Scale bars in panels A, D, F, G = 5 µm. (**B**) The AF2 model confidence score (pLDDT, displayed blue to red through green from highly to poorly reliable) highlights regions of the model predicted with high or poor confidence, respectively (Jumper et al., 2021; Tunyasuvunakool et al., 2021). (**C**) A predicted N-terminal extension (green) of the ROD *β*-propeller (red), slightly rotated from the view in B. The propeller proper begins with strand *β*7d and ends with strand *β*7c, which leads into the helical domain. The N-terminal extension augments the sixth and seventh blades with external *β*-strands (*β*6e and *β*7e). The position of T13 and S15 on the extension is shown. (**D**) The phosphorylation state of RZZ and Spindly^F^ was monitored by ProQ Diamond after SDS-PAGE separation of reactions. (**E**) Filamentation assays with the indicated combinations of 8 µM ^mCh^RZZS^F^, MPS1 (1 µM), and Lambda phosphatase (0.4 mg/ml) in presence of 10 µM reversine. Dephosphorylation reactions were carried out on already formed filaments (see Methods). Samples were imaged by confocal microscopy. Dephosphorylation does not dissolve already formed filaments. (**F**) An N-terminal deletion mutant of ROD (Δ15) removing the first 15 N-terminal residues that include the MPS1 phosphorylation sites Thr13 and Ser15 is unable to form filament in presence of MPS1 at 20°C, but can form filaments upon mildly heating to 30°C. (**G**) RZZ and RZZS complexes were reconstituted with the Zwilch^E422A/D426A^ mutant and tested for filamentation at 20°C in presence of MPS1 or upon mildly heating to 30°C. These experiments are also displayed in Figure 5 – Supplement 1A (**H**) Surface representation of the RZZ model depicting the position of Zwilch^E422^ and Zwilch^D422^ (in purple) and the positions of the ROD N-terminal region (green) and the highly conserved ZW10 N-terminus.

The inability of the T13A/S15A mutant of ^mCh^RZZS^F^ to form fibers did not reflect a requirement of these residues in fiber assembly, because fibers of this mutant were observed after mildly heating the sample to 30°C (Figure 5 – supplement 1A). Fibers never formed when Spindly^F^ was omitted (Figure 5 – supplement 1B). Thus, T13 and S15 may not be required for a direct physical interaction of ^mCh^RZZS^F^ complexes (protomers) in the fiber. Rather they may be required for auto-inhibition of filament nucleation and growth. To shed light on how this region of ROD controls fiber formation, we asked if we could structurally model it. The ROD *β*-propeller begins with the *β*7d strand around residue Leu30 (Figure 5B-C). The N-terminal residues that precede this point of entry into the *β*-propeller cannot be mapped with certainty in the reconstruction, due to its limited resolution in this peripheral region. However, AF2 predicts that this region first augments blade 7 of the ROD *β*-propeller with an additional *β*-strand (*β*7e), external to the outermost *β*7d of the 4-strand propeller; then, after forming a helical turn that packs against the top part of blade 7, it also augments blade 6 with a short *β*6e strand that pairs with *β*6d, before finally entering the *β*-propeller (Figure 5C).

Based on these observations, we speculate that phosphorylation of Thr13 and Ser15 by MPS1 restructures the N-terminal region of ROD, relieving auto-inhibition and allowing interactions required for the nucleation and growth of RZZS filaments, possibly mediated by blades 6 and 7. To test this idea, we reasoned that if MPS1 phosphorylation of ROD were exclusively required to promote nucleation of ^mCh^RZZS^F^ filaments, the stability of already formed filament should remain unaffected after ROD dephosphorylation. Indeed, successful dephosphorylation of ^mCh^RZZS^F^ filaments with lambda phosphatase (Figure 5D) did not visibly interfere with filament number or stability (Figure 5E). An ^mCh^RZZS^F^ complex containing a ^mCh^ROD deletion mutant lacking the N-terminal 15 residues (^mCh^ROD^Δ15^) was insensitive to MPS1 phosphorylation, but, like the T13A/S15A mutant, formed fibers when heat-treated (Figure 5F). Thus, ^mCh^ROD^Δ15^ remains auto-inhibited, probably due to the residual capping of blade 6 with the *β*6e strand, which we speculate to be crucially required for fiber assembly.

### Zwilch contributes directly to fiber assembly

Previous studies implicated two highly conserved Zwilch residues, Glu422 and Asp426, in corona expansion in humans and nematodes (Gama et al., 2017; Pereira et al., 2018). To assess if these residues also have a direct effect on corona assembly *in vitro*, we engineered a ^mCh^RZZS^F^ complex containing alanine mutants of these residues (E422A/D426A). The E422A/D426A mutant complex failed to form fibers, both upon MPS1 phosphorylation and upon heating (Figure 5G). This was not caused by an impairment of the interaction of RZZ with Spindly, as the latter was unperturbed (Figure 5 – Supplement 1C). Thus, Zwilch contributes directly to fiber assembly. The two conserved Zwilch residues are solvent exposed and are part of a continuous face of the RZZ complex that also comprises blade 6 of the ROD *β*-propeller and the N-terminal region of ZW10 (Figure 5H), suggestive of an extensive interaction interface for corona expansion.

### The RZZS polymers

To shed further light into the polymerization mechanism, we used negative-stain EM to visualize the ^mCh^RZZS^F^ fibers obtained after mild heating or incubation with MPS1 kinase. Under either condition, the fibers appeared as flat sheets, consisting of somewhat irregular filaments packing side-a-side. The sheets co-existing with unpolymerized complexes and with small oligomers (Figure 6A). Essentially identical sheets were obtained with a complex containing untagged ROD (RZZS^F^, Figure 6 – Supplement 1A), or with complexes expressing the ROD mutants T13A/S15A, T13E/S15E, and Δ15 (Figure 6 – Supplement 1B). The sheets were not sufficiently ordered for a successful application of cryo-EM reconstruction methods.

**Figure 6.**
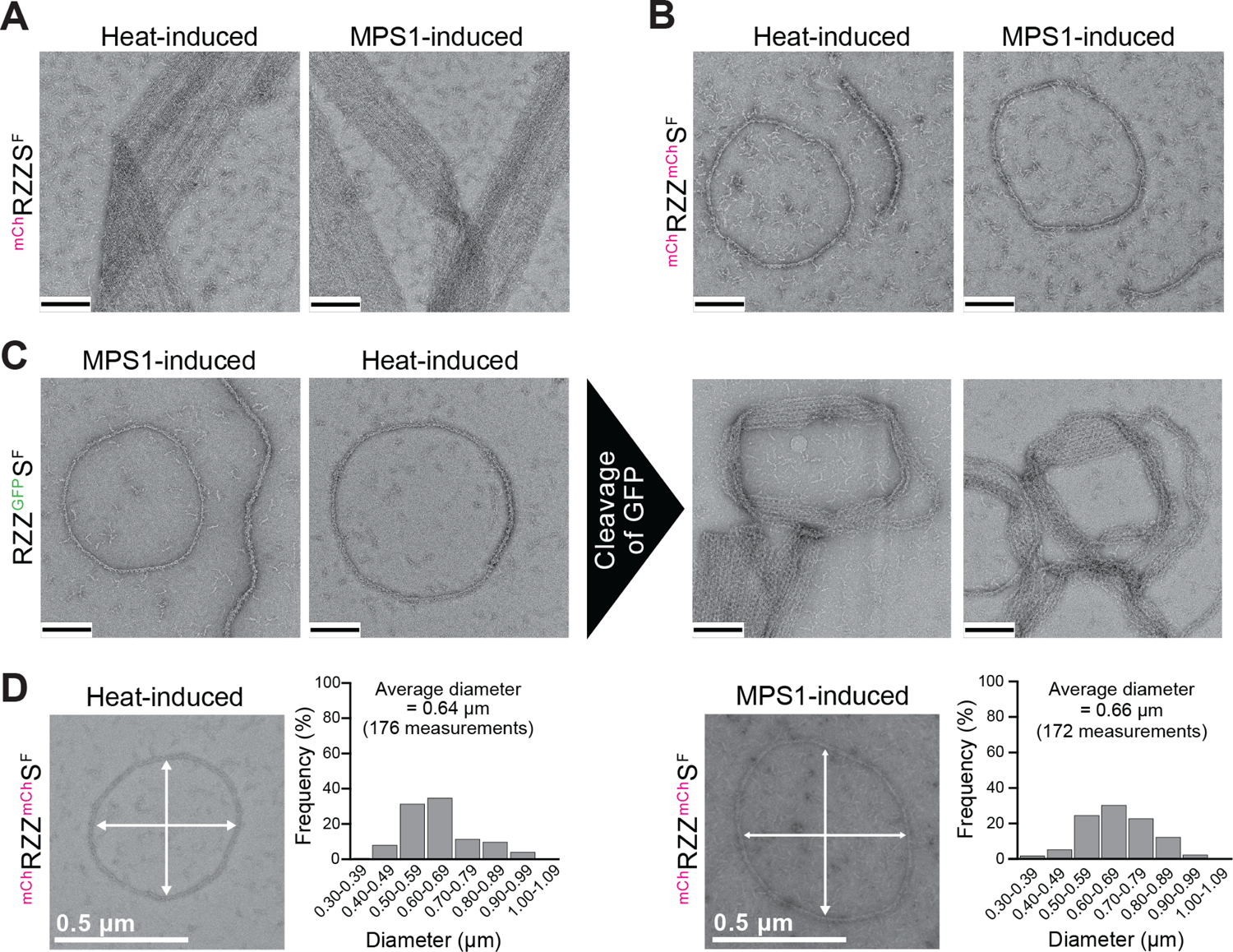
Ultrastructural analysis of RZZS sheets and filaments. (**A**) Negative-stain electron microscopy analysis of heat- and MPS1-induced sheets of filament of farnesylated ^mCh^RZZS. Scale bar (black): 200 nm. (**B**) ^mCh^RZZ^mCh^S forms rings and curved single filaments rather than sheets. Scale bar: 200 nm. (**C**) The GFP of Spindly was removed with Prescission protease, and the resulting objects were imaged by negative stain EM. Scale bar: 200 nm. (**D**) Heat- and MPS1-induced rings of ^mCh^RZZ^mCh^S have similar diameters.

Polymerization attempts with an RZZS complex where Spindly was also tagged with an N-terminal mCherry moiety (^mCh^RZZ/^mCh^S^F^) prevented formation of fibers and rather promoted formation of complete rings, or of segments thereof of comparable curvature (Figure 6B), and regardless of whether initiated by heat or MPS1. Essentially identical figures were also observed with an equivalent complex containing GFP-tagged Spindly (^mCh^RZZ/^GFP^S^F^; Figure 6 – Supplement 1C), even with untagged ROD (RZZ/^GFP^S^F^; Figure 6C). Cleavage of the GFP moiety from the latter construct after polymerization into rings promoted the lateral association of the filamentous rings into bundled rings, with a texture that was considerably less dense than that of the sheets, suggesting that bundles of rings do not pack as tightly as bundles of filaments in the sheets (Figure 6C).

Thus, N-terminal tagging of Spindly promotes the assembly of rings or curved filaments. The curvature of the rings, whose average diameter is approximately 0.65 µm (Figure 6D), is remarkably similar to the curvature of kinetochore crescents when the corona expands (Magidson et al., 2015). Two-dimensional (2D) class averages of short segments of the negatively stained samples comprising a few consecutive ring subunits revealed a substantial orientation preference that ultimately prevented the successful calculation of a 3D reconstruction (Figure 6 – Supplement 1D). Similar analyses on filaments at cryogenic temperatures suffered from the same extreme orientation preferences and were unsuitable for coherent reconstructions (unpublished data). Nonetheless, these analyses revealed that the rings appear to have a period of approximately half of the RZZ length (*≍*23 nm) and a width comparable to that of the RZZ (*≍*11 nm) (Figure 6 – Supplement 1E), suggesting that they form through staggering of individual RZZS complexes.

### The role of Spindly

To shed light on how Spindly promotes corona assembly, we investigated the corona assembly propensity of various Spindly deletion mutants, including Spindly^250-C^ and Spindly^354-C^, where the CC1a/b segment of Spindly (250-C) or the CC1a/b and CC2 segments (354-C) are deleted, respectively (Figure 3A). Both Spindly constructs, upon farnesylation, interacted with the RZZ complex in size-exclusion chromatography experiments (Figure 7A). Conversely, a further deletion construct, Spindly^440-C^, was unable to interact with the RZZ complex.

**Figure 7.**
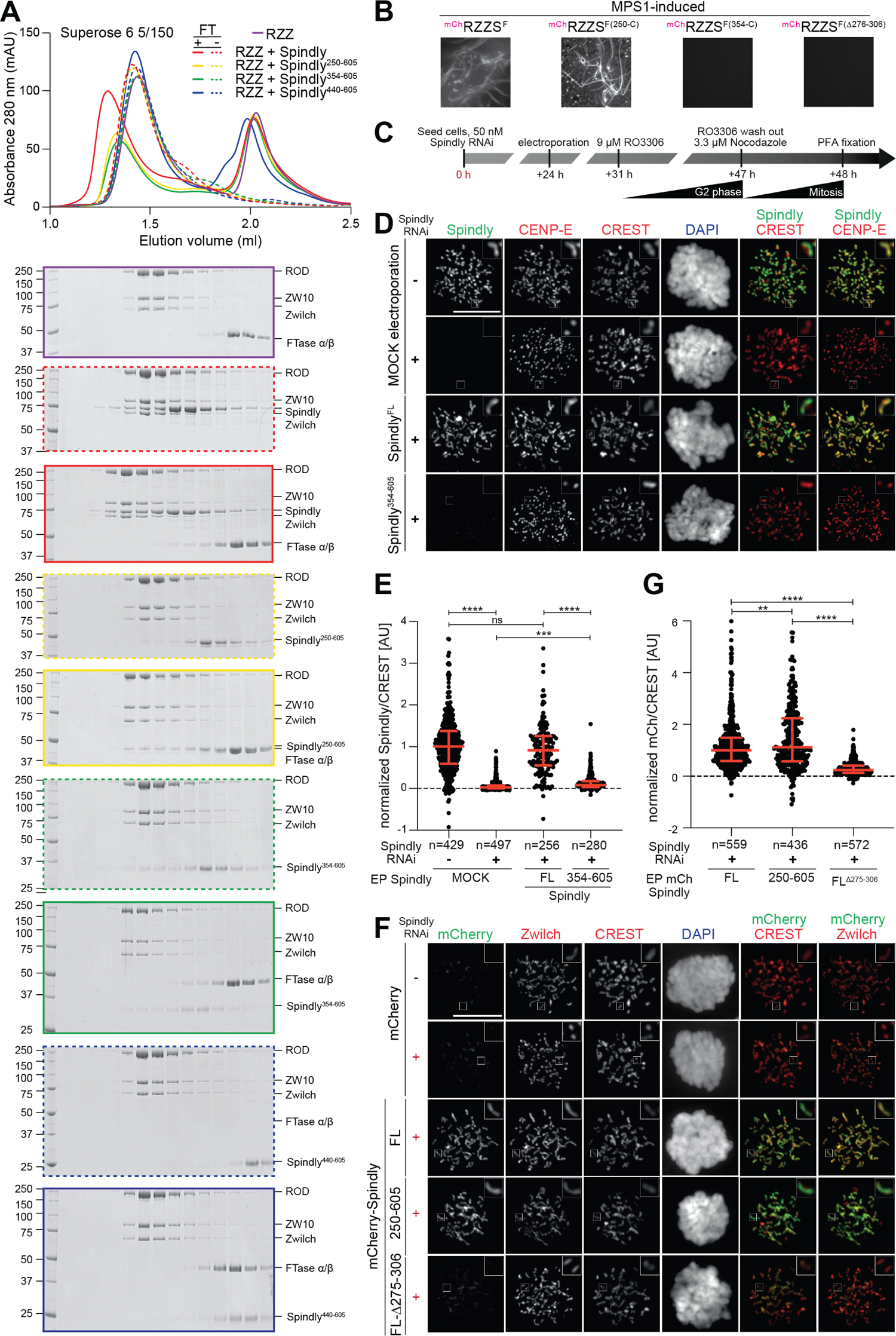
Influence of Spindly on corona assembly and kinetochore recruitment. (**A**) Size-exclusion chromatography and SDS-PAGE of elution fractions of the indicated samples. Each Spindly construct was incubated with RZZ in absence (dotted lines) or in presence (continuous lines) of farnesyl transferase and farnesyl pyrophosphate. Elution shifts of Spindly^F^ and of the resulting RZZS^F^ complexes is indicative of successful interaction. (**B**) MPS1-induced filamentation experiments on the indicated ^mCh^RZZS^F^ complexes (4 µM, further diluted to 0.5 µM for imaging) using a confocal spinning disk fluorescence microscope at 561 nm. (**C**) Schematic of the cell synchronization and imaging experiment shown in D. After electroporation, cells were allowed to recover for 8 hours. Subsequently, cells were synchronized in G2 phase with 9 µM RO3306 for 15 hours and then released into mitosis by inhibitor washout. Before fixation, cells were treated with 3.3 µM nocodazole for 1 hour. (**D**) Representative images of fixed HeLa cell electroporated with full-length Spindly and Spindly^354-605^ constructs in cells depleted of endogenous Spindly by RNAi. Spindly localization was detected with an antibody against the C-terminal region of Spindly. Corona expansion or lack thereof were monitored through CENP-E. CREST serum was used to visualize kinetochores, DAPI to stain for DNA. Scale bar: 10 µm. (**E**) Scatter dot plots representing normalized intensity ratios of the indicated Spindly constructs over CREST for individual kinetochores of cells from the experiment shown in panel **D**. Red lines indicate median with interquartile range. (**F**) Cells treated like in C-D were electroporated with the indicated ^mCh^Spindly constructs. Corona expansion was evaluated through the appearance of Zwilch. (**G**) Kinetochore intensities were quantified like in panel E.

While Spindly^250-C^ supported corona expansion *in vitro* upon MPS1 phosphorylation and mild heating, indistinguishably from full-length Spindly, Spindly^354-C^ did not support corona assembly (Figure 7B and Figure 7 – Supplement 1A). Furthermore, while Spindly^354-C^ bound the RZZ *in vitro*, it was unable to decorate kinetochores after introduction by electroporation into HeLa cells depleted of endogenous Spindly (Figure 7C-D, quantified in panel E). As expected, therefore, there was no expansion of the corona in these cells, as shown by the dot-like appearance of CENP-E, contrasting its crescent-like appearance observed in control cells with a well-formed corona. Spindly decorates kinetochores even when corona expansion is suppressed with an MPS1 inhibitor (Figure 4E), indicating that its localization to kinetochores is not contingent on corona expansion. Thus, failure of Spindly^354-C^ to reach kinetochores is unlikely to reflect its inability to assemble the corona (Figure 7C). Rather, Spindly^354-C^ may bind RZZ with reduced affinity, or may be unable to interact with one or more additional kinetochore receptors ultimately required to stabilize the RZZS complex.

Collectively, these results indicate that a segment of Spindly encompassing residues 250-353 contains a critical determinant of corona assembly and kinetochore recruitment, possibly distinct or even overlapping. To shed further light on this question, we took advantage of our previous observation that Spindly^1-275^ and Spindly^306-C^ are stable Spindly construct (Mosalaganti et al., 2017; Sacristan et al., 2018) to build a new deletion mutant, Spindly^Δ276-306^. Spindly^Δ276-306^ bound robustly to the RZZ complex in size-exclusion chromatography experiments (Figure 7 – Supplement 1B), but was unable to support corona expansion *in vitro* (Figure 7B) and failed to reach the kinetochore (Figure 7F, quantified in panel G; note that the very modest mCherry signal shown to overlap with the centromere and inner kinetochore is a localization artifact of the mCherry tag). In conclusion, these results identify a segment comprising 31 residues of Spindly (276-306) as a crucial determinant of Spindly kinetochore localization and corona expansion.

## Discussion

The high-resolution cryo-EM structure of the RZZ complex reported here crowns a succession of studies that began with the biochemical reconstitution and bioinformatic analysis of RZZ subunits, the determination of the crystal structure of Zwilch, and the report of a low-resolution EM reconstruction of the RZZ (Altenfeld et al., 2015; Civril et al., 2010; Mosalaganti et al., 2017). Our new structural analysis leveraged a pipeline that combined experimental structure determination using cryo-EM, model building based on experimental 3D reconstructions, and the enhanced prediction capabilities of AlphaFold2 (Jumper et al., 2021; Tunyasuvunakool et al., 2021), which were instrumental for model building in more peripheral regions of the reconstruction where the local resolution did not allow *de novo* model building. Collectively, this pipeline allowed us to reveal the structure of the RZZ complex at near-atomic resolution. The new structure explains how intermolecular interactions of the subunits promote the assembly of the RZZ complex; it also explains how RZZ interacts with the farnesyl moiety of Spindly; finally, it sets the basis for understanding how RZZ assembles into supramolecular structures in the corona.

We refined an *in vitro* assay for corona reconstitution that allowed us to probe several aspects of the polymerization reaction. First, we demonstrate that RZZS oligomerization *in vitro* into flat sheets or rings is kinetically controlled, and can be induced either by raising temperature or by addition of the MPS1 kinase. Under all tested conditions, RZZ polymerization *in vitro* required Spindly^F^, implicating it as a crucial building block of the corona. Two non-mutually exclusive possibilities are that Spindly^F^ contributes directly to binding interfaces required for polymerization, or that it induces a conformational change in the RZZ required for polymerization. Our initial efforts to reveal the structure of the RZZS^F^ filament were thwarted by the limited order of the fibers we have obtained and by a very limited number of orientations on the EM grids. Future work will have to address this bottleneck, shedding light on the organization of the individual RZZS complex and of its polymeric form.

Nonetheless, our studies identified and mutationally probed several crucial interfaces for polymerization, including the N-terminal region of Spindly and a conserved acidic residue pair in Zwilch. Human RZZS^F^ polymerizes efficiently at 30°C *in vitro* in the absence of a kinetochore support, whereas its polymerization in cells is seeded by the kinetochore and never extends far from it. While this may seem to suggest that other control mechanisms prevent RZZS oligomerization away from kinetochores, it should be considered that our experiments *in vitro* were carried out at RZZS^F^ concentrations (usually 4 µM) likely to be approximately two orders of magnitude higher than those existing in cells, as inferred by the fact that most SAC components have concentrations comprised between 10 and 100 nM (Simonetta et al., 2009). High concentration of building blocks likely facilitates polymerization, and indeed RZZS^F^ filaments became sporadic or were not any longer observed at mid-nanomolar concentrations of RZZS^F^ (Figure 7 – Supplement 1C). A second crucial factor likely explaining why RZZS filaments form only at kinetochores is that RZZS polymerization appears to be kinetically controlled, with MPS1 phosphorylation acting as catalyst to remove a steric blockade to oligomerization involving the ROD N-terminal region. As kinetochores enrich MPS1 during mitosis, and MPS1 activity is highest at these structures, albeit not limited to them (Kuijt et al., 2020), polymerization may become naturally spatially limited to kinetochores.

This mechanism of corona assembly bears similarities to the process of coat assembly that drives intracellular trafficking of membranous organelles. In addition to evident structural similarities, most notably of ROD with Clathrin and COPs, which are also characterized by a succession of an N-terminal *β*-propeller and a C-terminal *α*-solenoid, both processes are spatially and kinetically controlled so that they occur only at defined cellular locales and in presence of appropriate triggers (Arakel and Schwappach, 2018; Sigismund et al., 2021). The high-affinity binding site that drives RZZ recruitment to the kinetochore remains elusive, but appears to be confined within the KMN network (Caldas et al., 2015; Chan et al., 2009; Miller et al., 2008; Pagliuca et al., 2009; Pereira et al., 2018; Sundin et al., 2011; Varma et al., 2013).

At least two kinases, in addition to MPS1, are also required for assembly and/or retention of the corona, CDK1 and Aurora B. Acute inhibition of these kinases results respectively in the detachment of the assembled corona from the kinetochore (CDK1) and in the complete depletion of corona components at kinetochores (Aurora B). In experiments with purified kinases *in vitro*, we did not find prominent Aurora B phosphorylation sites on the RZZS^F^ complex, suggesting that Aurora B does not controls corona assembly directly. Because Aurora B is critically required for MPS1 recruitment to kinetochores (Nijenhuis et al., 2013; Santaguida et al., 2010), we suspect that Aurora B inhibition blocks the essential function of MPS1 as promoter of corona assembly. On the other hand, CDK1 phosphorylates Spindly^F^ efficiently *in vitro*, but without triggering filamentation. It is possible that the detachment of the corona after CDK1 inhibition reflects an essential role of CDK1 phosphorylation of Spindly in its kinetochore recruitment (e.g. by creating a phospho-dependent binding site or conformational change). In this case, CDK1 inhibition may be recapitulated by expression of non-phosphorylatable mutants of Spindly. Alternatively, CDK1 may contribute to the generation of a binding site for the RZZS on the kinetochore.

Previous observations identified RZZ and Spindly as being both necessary for corona assembly in human cells (Rodriguez-Rodriguez et al., 2018; Sacristan et al., 2018). Our results *in vivo* are consistent with this tenet, but are further supported by polymerization experiments *in vitro* that showed a nearly perfect correlation with corona assembly in living cells. This coincidence argues that the determinants of corona assembly, after excluding the unknown receptor site in the kinetochore, are entirely contained with the RZZS complex. Thus, the RZZS emerges from our study as being sufficient to assemble the corona. This conclusion also explains why depletion of additional corona components, including CENP-E, CENP-F, DD, and MAD1-MAD2, does not visibly disrupt corona assembly and RZZ kinetochore recruitment (Allan et al., 2020; Ciossani et al., 2018; Gassmann et al., 2010). These proteins, on the other hand, may retain residual kinetochore localization even after depletion of RZZS components. The MAD1-MAD2 complex, for instance, requires the RZZ complex for kinetochore localization, and will localize to the kinetochore even if the corona cannot expand due to Spindly depletion (Rodriguez-Rodriguez et al., 2018). In another example, CENP-E and CENP-F, in addition to interacting with the corona, have also been shown to interact with the non-corona components BUBR1 and BUB1, respectively (Ciossani et al., 2018; Legal et al., 2020; Raaijmakers et al., 2018). We anticipate that our corona assembly assay may shed light on the mechanism of recruitment of these additional corona components.

A fundamental unresolved question in kinetochore biology is how the conversion of microtubule attachments from lateral to end-on promotes corona stripping. Plausibly, this sudden transition reflects a weakening of the interaction of the RZZS with its kinetochore receptor, leading to DD-directed detachment of the corona from the kinetochore. What triggers this change in binding affinity, however, remains unclear. Our studies, by unveiling the molecular features of the RZZ complex and by defining requirements for its physical interactions, provide an initial step towards the elucidation of this very complex and important question.

## Acknowledgments

We are grateful to Oliver Hofnagel and Daniel Prumbaum for help in EM data collection, and to the Musacchio and Raunser laboratories for helpful discussion. This work was supported by the Max Planck Society (to A.M. and S.R.). A.M. acknowledges funding by the Marie-Curie Training Network DivIDE (project number 675737), European Research Council (ERC) through Synergy Grant 951439 (BIOMECANET), the Deutsche Forschungsgemeinschaft (DFG, German Research Foundation) through SFB1430 (Project-ID 424228829). Density maps of RZZ will be deposited into EMDB upon publication. Coordinates of the RZZ complex will be deposited to the PDB upon publication. The authors declare no competing financial interests.

## Author contributions (following CRediT model) Conceptualization

**Andrea Musacchio**

**Data curation:** N/A

**Formal analysis:** Tobias Raisch, Ingrid Vetter, Felipe Merino

**Funding acquisition:** Andrea Musacchio, Stefan Raunser

**Investigation:** Verena Cmentowski, Giuseppe Ciossani, Ennio d’Amico, Tobias Raisch

**Project Administration:** Andrea Musacchio, Stefan Raunser

**Resources:** Sara Carmignani, Stefano Maffini, Sabine Wohlgemuth

**Supervision:** Stefano Maffini, Andrea Musacchio, Stefan Raunser, Ingrid Vetter

**Validation:** Andrea Musacchio, Stefan Raunser, Ingrid Vetter

**Visualization:** Verena Cmentowski, Giuseppe Ciossani, Ennio d’Amico, Andrea Musacchio, Tobias Raisch, Ingrid Vetter

**Writing – original draft:** Andrea Musacchio

**Writing – review & editing:** Sara Carmignani, Giuseppe Ciossani, Verena Cmentowski, Ennio d’Amico, Stefano Maffini, Felipe Merino, Andrea Musacchio, Tobias Raisch, Stefan Raunser, Ingrid Vetter

## Materials and Methods

### Protein expression and purification

Expression and purification of Spindly and RZZ constructs were carried out essentially as previously described (Mosalaganti et al., 2017; Sacristan et al., 2018). Proteins were expressed using the BiGBac recombinant expression system (Weissmann et al., 2016). Bacmid was produced from EMBacY cells and used to transfect Sf9 cells to produce baculovirus. The baculovirus was subjected to three rounds of amplification, and used to infect TnaO38 cells. Cells were cultured for 72 hours before harvesting. A pellet from expression in 500 ml of TnaO38 cells was lysed by sonication in 100 ml lysis buffer (50 mM HEPES pH 8.0, 250 mM NaCl, 2 mM TCEP, 50 mM imidazole) supplemented with 1 mM PMSF and protease inhibitor cocktail (Serva). The lysate was then cleared by centrifugation at 100000 g for 45 minutes. The cleared lysate was loaded onto a HisPrep Fast Flow column (Cytiva) pre-equilibrated in lysis buffer, and washed with 500 ml lysis buffer, after which the bound protein was eluted with lysis buffer supplemented with 300 mM imidazole. The eluate was diluted 1:5 in Buffer A (50 mM HEPES pH 8.0, 2 mM TCEP), and applied to a 6 ml Resource Q anion exchange column (Cytiva). The protein was then eluted using a 50 mM to 500 mM NaCl gradient. Peak fractions were analyzed by SDS-PAGE, and those containing the relevant proteins were concentrated to 500 µl volume, and subjected to size-exclusion chromatography on a Superdex S200 10/300 column (Cytiva) for Spindly, and on a Superose 6 10/300 for RZZ constructs, equilibrated in SEC buffer (50 mM HEPES pH 8.0, 250 mM NaCl, 2 mM TCEP). Fractions were pooled and concentrated to 10 mg/ml, snap-frozen, and stored at −80 °C until use. For dephosphorylation, 4 µM ^mCh^RZZS^F^ complex or pre-formed MPS1-induced ^mCh^RZZS^F^ polymers were incubated for 15 hours in M-buffer at 20°C with or without 0.4 mg/ml Lambda phosphatase (produced in house), in presence of 10 µM reversine.

Spindly^Δ276-306^ was expressed as an mCherry fusion in E. coli. BL21 CodonPlus cells were transformed with a pET28a plasmid containing the coding sequence for the mCherry tag, a PreScission cleavage sequence, and Spindly^Δ276-306^, and grown in TB at 37°C to an OD_600_ of 0.5. Expression was induced with 0.4 mM IPTG. The culture was then transferred into an incubator pre-cooled to 18° C, and grown overnight before harvesting. The pellet was then snap-frozen and stored at −80° C until purification. Untagged Spindly^Δ276-306^ was obtained by cleaving the mCherry tag with PreScission protease, by incubating 2 mg of mCherry-Spindly^Δ276-306^ with 0.1 mg of in-house produced PreScission protease overnight. The sample was then loaded on a Superose 6 column equilibrated in SEC buffer to remove the tag and PreScission protease. Fractions were then pooled and concentrated to 10 mg/ml, snap-frozen, and stored at −80° C until use.

### Production of MPS1 kinase

An mCherry-MPS1-6His construct was generated by sub-cloning in a pLIB vector. The corresponding baculovirus was generated in Sf9 insect cells (Wickham et al., 1992). After three rounds of amplification (V0, V1, and V2, 4 days each), 100 ml of V2 were inoculated in 1 liter of Tnao38 cells. 24 hours after infection for expression, reversine (Santaguida et al., 2010) was added to the growth medium (1 µM) to maximize expression yields. After 60 hours of expression at 27°C, cells were pelleted, washed in PBS, pelleted again and either stored at −80°C after flash-freezing in liquid nitrogen, or used directly for purification. Every purification step was performed on ice or at 4°C. The pellet was resuspended in ∼100 ml buffer A (300 mM NaCl, 50 mM Hepes pH 8, 5% Glycerol, 2 mM TCEP, 10 mM Imidazole pH 8) and supplemented with PMSF (1:100), protease-inhibitor mix HP Plus (1:500, Serva) and DNaseI (1:300, Roche), lysed by sonication and cleared by centrifugation at 108000g for 45 min. The cleared lysate was applied to 5 ml Nickel-NTA (GE Healthcare) slurry beads previously equilibrated in buffer A and incubated on a rotating platform at 4°C for 2 hours. The supernatant was removed by centrifugation (1500g, 5 min, 4°C) and the beads were washed with 100 ml buffer A. For the elution, the beads were incubated (∼15 min at 4°C) in 15 ml of buffer A supplemented with 300 mM Imidazole pH 8. Samples of the cleared lysate, of the supernatant, and of the elution were loaded on SDS-page for analysis. The 15 ml elution was then concentrated, spun at max speed for 30 min in a bench-top centrifuge (at 4°C) and finally loaded on a Superdex200 16/60 (GE Healthcare) previously equilibrated in MPS1 buffer (250 mM NaCl, 50 mM Hepes pH 8, 5% glycerol, 2 mM TCEP). The protein was then concentrated, aliquoted and stored in −80°C after flash-freezing in liquid nitrogen.

### In vitro farnesylation

Farnesyltransferase *α*/*β* mutant (W102T/Y154T) was expressed and purified as previously described (Mosalaganti et al., 2017). Spindly was diluted to 100 µM in farnesylation buffer (50 mM HEPES pH 8.0, 250 mM NaCl, 10 mM MgCl2, 2 mM TCEP), and farnesyltransferase was added to a final concentration of 30 µM. Farnesyl pyrophosphate (Sigma-Aldrich) was added stepwise to a final concentration of 300 µM. The reaction mixture was incubated at RT for 6 hours, after which it was centrifuged at 16,000 g for 10 minutes to remove precipitate that formed during the reaction. The cleared reaction mixture was then loaded on a Superose 6 column equilibrated in SEC buffer to remove the farnesyltransferase. The fractions containing Spindly were identified by SDS-PAGE and pooled, concentrated to a final concentration of around 5 mg/ml, snap-frozen, and stored at −80 °C until use.

### Analytical size-exclusion chromatography

Analytical gel filtration runs were performed on Superose 6 Increase 5/150 columns (Cytiva) pre-equilibrated in SEC buffer. For runs with pre-farnesylated Spindly, RZZ and Spindly were pre-incubated at a concentration of 2 µM and 6 µM respectively on ice for 1 h in SEC buffer. For runs with concurrent farnesylation, RZZ, Spindly, and FTase were incubated at a concentration of 5 µM, 15 µM and 7.5 µM respectively for 2 h at room temperature in SEC buffer supplemented with 25 µM FPP, followed by 30 min on ice.

### Filamentation experiments with RZZS or RZZ complexes

Heat-induced RZZS^F^ filaments were prepared by incubating 4 µM RZZ complex and 8 µM Spindly^F^ or 4 µM preassembled RZZS^F^ complex for 1 h at 30°C in H-buffer (50 mM Hepes pH 7.5, 100 mM NaCl, 1 mM MgCl_2_ and 1 mM TCEP). Tags were removed from ^mCherry^RZZ^GFP^S^F^ filaments by incubating the polymers for 1 hour at 30°C with 0.5 mg/ml Prescission protease (produced in house). MPS1-induced filaments were obtained by incubating 4 µM RZZ complex and 8 µM Spindly^F^ or 4 µM RZZS^F^ preassembled purified complex for 15 hours at 20°C in M-buffer (50 mM Hepes pH 7.5, 100 mM NaCl, 1 mM MgCl_2_ and 1 mM TCEP), supplemented with 2 mM ATP at pH 8.0 and in presence of 1 µM MPS1. The effect of other mitotic kinases on RZZS^F^ complex filamentation was tested in the same conditions using 1 µM of purified protein kinase (produced in house). Reversine (Calbiochem) was dissolved at 10 mM in DMSO and used at 10 µM final concentration. Protein phosphorylation was monitored by ProQ Diamond phosphostaining (ThermoFisher Scientific) after SDS-PAGE separation. Independent MPS1 phosphorylation of RZZ and Spindly^F^ protein stocks was carried out by incubating 8 µM RZZ and 16 µM Spindly^F^ overnight in M-buffer at 20°C, supplemented with 2 mM ATP pH 8.0 and 1 µM MPS1.

### Confocal imaging of RZZS^F^ filaments

Glass flow chambers of about 10 µl volume were assembled using standard cover glasses and glass slides, held together by double-side tape (Teva). Heat- or MPS1-induced RZZS^F^ filaments were diluted to 0.5 µM in H- or M-buffer (see above), respectively, and imaged in the glass chambers, at room temperature using a spinning disk confocal device on the 3i Marianas system at 63X magnification. Sample images were acquired as 5-stacks of z-sections at 0.27 µm, converted into maximal intensity projections, exported and processed with Fiji (Schindelin et al., 2012).

### Cell culture, plasmid transfection, microinjections and imaging

<u>Cell culture and drug treatment:</u> HeLa and DLD-1 cells were grown in Dulbecco’s Modified Eagle’s Medium (DMEM; PAN Biotech) supplemented with 10 % tetracycline-free FBS (PAN Biotech), and L-glutamine (PAN Biotech). Cells were grown at 37°C in the presence of 5 % CO_2_. Where indicated, nocodazole (Sigma) was used at 3.3 µM, RO3306 (Calbiochem) at 9 µM, MG-132 (Calbiochem) at 10 µM, hesperadin at 500 nM (Merck), and reversine (Cayman Chem.) at 500 nM. <u>Cell transfection and electroporation:</u> Depletion of endogenous Spindly was achieved through reverse transfection with 50 nM Spindly siRNA (5′-GAAAGGGUCUCAAACUGAA-3′ obtained from Sigma-Aldrich) for 48 hours with RNAiMAX (Invitrogen, Carlsbad, California, United States). For rescue experiments, 24 hours after Spindly depletion, we electroporated recombinant Spindly constructs, either unlabeled or labeled with an N-terminal mCherry, at a concentration of 7 μM in the electroporation slurry (as previously described in Alex et al., 2019) (Neon Transfection System, Thermo Fisher). Control cells were electroporated with mCherry or electroporation buffer, respectively. Following an 8 hours recovery, cells were treated with 9 µM RO3306 (Calbiochem) for 15 hours. Subsequently, cells were released into mitosis in presence of 3.3 µM nocodazole (Sigma) for 1 hour. <u>Immunofluorescence:</u> Cells were grown on coverslips pre-coated with Poly-L-lysine (Sigma-Aldrich). Cells were pre-permealized with 0.5% Triton X-100 solution in PHEM (Pipes, HEPES, EGTA, MgCl_2_) buffer supplemented with 100 nM Microcystin for 5 minutes before fixation with 4% PFA in PHEM for 20 minutes. After blocking with 5% boiled goat serum (BGS) in PHEM buffer, cells were incubated for 2 hours at room temperature with the following primary antibodies: BUB1 (mouse, Abcam, ab54893, 1:400), CENP-E (mouse, Abcam, ab5093, 1:200), Spindly (rabbit, Bethyl, A301-354A, 1:1000), Zwilch (rabbit, made in-house, SI520, 1:900), CREST/anti-centromere antibodies (Antibodies, Inc., 1:200) diluted in 2.5 % BGS-PHEM supplemented with 0.1% Triton-X100. Subsequently, cells were incubated for 1 hour at room temperature with the following secondary antibodies: Goat anti-mouse Alexa Fluor 488 (Invitrogen A A11001), donkey anti-rabbit Rhodamine Red (Jackson Immuno Research 711-295-152), donkey anti-rabbit Alexa Fluor 488 (Invitrogen A21206), goat anti-human Alexa Fluor 647 (Invitrogen, Carlsbad, California, United States). All washing steps were performed with PHEM-T buffer. DNA was stained with 0.5 μg/ml DAPI (Serva) and Mowiol (Calbiochem) was used as mounting media. <u>Cell imaging:</u> Cells were imaged at room temperature using a spinning disk confocal device on the 3i Marianas system equipped with an Axio Observer Z1 microscope (Zeiss), a CSU-X1 confocal scanner unit (Yokogawa Electric Corporation, Tokyo, Japan), 100 × /1.4NA Oil Objectives (Zeiss), and Orca Flash 4.0 sCMOS Camera (Hamamatsu). Images were acquired as z sections at 0.27 μm. Images were converted into maximal intensity projections, exported, and converted into 8-bit tiff files. Automatic quantification of single kinetochore signals was performed using the software Fiji with background subtraction. Measurements were exported in Excel (Microsoft) and graphed with GraphPad Prism 9.0 (GraphPad Software). The figures were arranged using Adobe Illustrator 2022 software.

### Negative stain electron microscopy sample preparation and image analysis

4 µl of 20-100 nM RZZS^F^ filaments were deposited on freshly glow-discharged Formvar/Carbon (Quantifoil) film supported copper grid Cu400 (Sigma Aldrich) and incubated for 1 min. Once removal of the excess of sample was blotted away with filter paper, the grids were washed two times with 10 µl of H- or M-buffer (Heat- or Mps1-induced filaments, respectively), then once with 10 µl of 0.75% (w/v) uranyl formate (Sigma Aldrich). After staining with 10 µl of 0.75% (w/v) uranyl formate for 30 sec, grids were blotted, dried and visualized at 120 kV using a Tecnai Spirit equipped with a LaB_6_ cathode and a 4000 × 4000 CMOS detector F416 (TVIPS). Images were recorded at a nominal magnification of 21-42,000x. Single measurements of the diameter of RZZS^F^ circular polymers were performed by processing negative stain EM images with Fiji (NIH). Values were exported in Excel (Microsoft) and graphed with GraphPad Prism 6.0 (GraphPad Software). 2D classification of ^mCh^RZZ^GFP^S^F^ filaments was performed using ISAC (Yang et al., 2012) within SPHIRE (Moriya et al., 2017). 148 images were collected at a magnification of 42000x resulting in 2.6 A/pix. Straight filament sections were manually selected, and segments of 256×256 px and an overlap of 115 px were extracted from those, resulting in 2730 particles. Classification was performed with standard parameters, using a radius of 120 px and a maximum of 50 members per class.

### Cryo-EM grid preparation and data acquisition

Grids were prepared using a Vitrobot Mark IV (Thermo Fisher Scientific) at 13 °C and 100 % humidity. 4 µl of RZZ at a concentration of 5 mg/ml and supplemented with 0.02 % Triton were applied to glow-discharged Quantifoil R2/1 grids and excess liquid removed by blotting (3.5 seconds at blot force −3) before vitrification in liquid ethane. Cryo-EM data were acquired on a Titan Krios electron microscope (Thermo Fisher Scientific) equipped with a field emission gun. Two datasets with 1968 and 5794 movies, respectively, were recorded on a K3 camera (Gatan) operated in super-resolution mode at a nominal magnification of 105,000, resulting in a super-resolution pixel size of 0.45 Å. A Bioquantum post-column energy filter (Gatan) was used for zero-loss filtration with an energy width of 15 eV. Total electron exposure of 59 and 60 e-/ Å2, respectively, was distributed over 60 frames. Data were collected using the automated data collection software EPU (Thermo Fisher Scientific), with three exposures per hole and a set defocus range of −1.2 to −2.7 µm. Details of data acquisition parameters can be found in Supplementary Table 1.

### Cryo-EM data processing

On-the-fly data pre-processing, including correction of beam-induced motion and dose-weighting by MotionCor2 (Zheng et al., 2017), CTF parameter estimation using CTFFIND4 in movie mode (Rohou and Grigorieff, 2015), and particle picking using a custom neural network in SPHIRE-crYOLO (Wagner et al., 2019), was performed within TranSPHIRE (Stabrin et al., 2020). 74,836 four-fold binned particles with dimensions of 180×180 pixels were extracted from the first dataset using SPHIRE (Moriya et al., 2017), and used for 2D classification in ISAC. An initial model was calculated in RVIPER using 81 good 2D classes and by imposing C2 symmetry. Initial 3D reconstruction in MERIDIEN was performed using 49,656 two-fold binned particles of the second dataset which were assigned to well-defined 2D classes and also with imposed C2 symmetry, resulting in a 4.7 Å 3D reconstruction. Recentered particles from all micrographs of the second dataset were used for training an improved neural network for SPHIRE-crYOLO. 191,979 particles were picked with this network on 7,718 micrographs of both datasets and extracted with two-fold binning and a size of 300×300 pixels. An initial 3D refinement of this particle stack in RELION (Zivanov et al., 2019) resulted in a 4.9 Å reconstruction. After several rounds of particle polishing and CTF refinement in RELION, reconstructions with overall nominal resolutions of 4.1 Å and 3.9 Å were obtained using 3D refinement in RELION or non-uniform refinement in cryoSPARC (Punjani et al., 2017; Punjani et al., 2020), respectively (Figure 1 – Supplement 1A-B). Local resolution was calculated, and the reconstruction locally filtered, using cryoSPARC (Figure 1 – Supplement 1C). As the global reconstructions displayed a strong resolution gradient from the center to the exterior parts of the molecule (Figure 1 – Supplement 1C), indicative of continuous flexibility within the complex, we turned to a focused refinement strategy. For this, we generated two focused masks, one for the central and one for the exterior part of one asymmetric half of the molecule. Then, we symmetry-expanded the particle stack according to the C2 symmetry and performed focused local refinements in cryoSPARC that resulted in reconstructions of 3.7 Å and 4.8 Å, respectively, for the central and the exterior masks (Figure 1 – Supplement 1A). The maps were fitted to the original unmasked C2-symmetric map, and a composite map created using the ‘vop maximum’ command in Chimera (Pettersen et al., 2004).

### Model building, validation, fitting

The central part of RZZ was built de novo using the 3.9 Å reconstruction from cryoSPARC non-uniform refinement. This map was subjected to automated model building using phenix.map_to_model (Liebschner et al., 2019). The resulting initial model, which comprised many α-helices in the central part of the particle, was manually improved and extended and the correct sequence assigned in Coot (Emsley et al., 2010), yielding a model comprising almost full-length ZW10 and the central part of Rod (residues 890-1440). AlphaFold2 (AF2) (Jumper et al., 2021; Tunyasuvunakool et al., 2021) was used in the original implementation as well as in the modified “ColabFold” (Mirdita et al., 2021) and in the “multimer” versions (Evans et al., 2021) to model subcomplexes of RZZ, specifically, the central region of ROD with ZW10, the “hook”-region consisting of the N- and C-termini of two different ROD molecules, the ROD *β*-propeller-Zwilch complex and the ROD *β*-propeller complex with the C-terminus of Spindly. The overlapping subcomplexes were superimposed, rigid-body-fitted to the RZZ electron density map, and then manually optimized. Flexible dynamic molecular dynamics fitting (Kidmose et al., 2019) and PHENIX real space refinement (Afonine et al., 2013) was employed to refine the fit and optimize the model geometries. In almost all regions, the AF2 predictions explained the electron density map very well after minor alterations, but the choice of the lengths of the interacting fragments of ROD was important, as e.g. the termini of full length ROD are predicted to interact with themselves since the second ROD molecule is missing, and a prediction of the dimeric full length complex was not successful.

---

**Supplementary Table 1.**
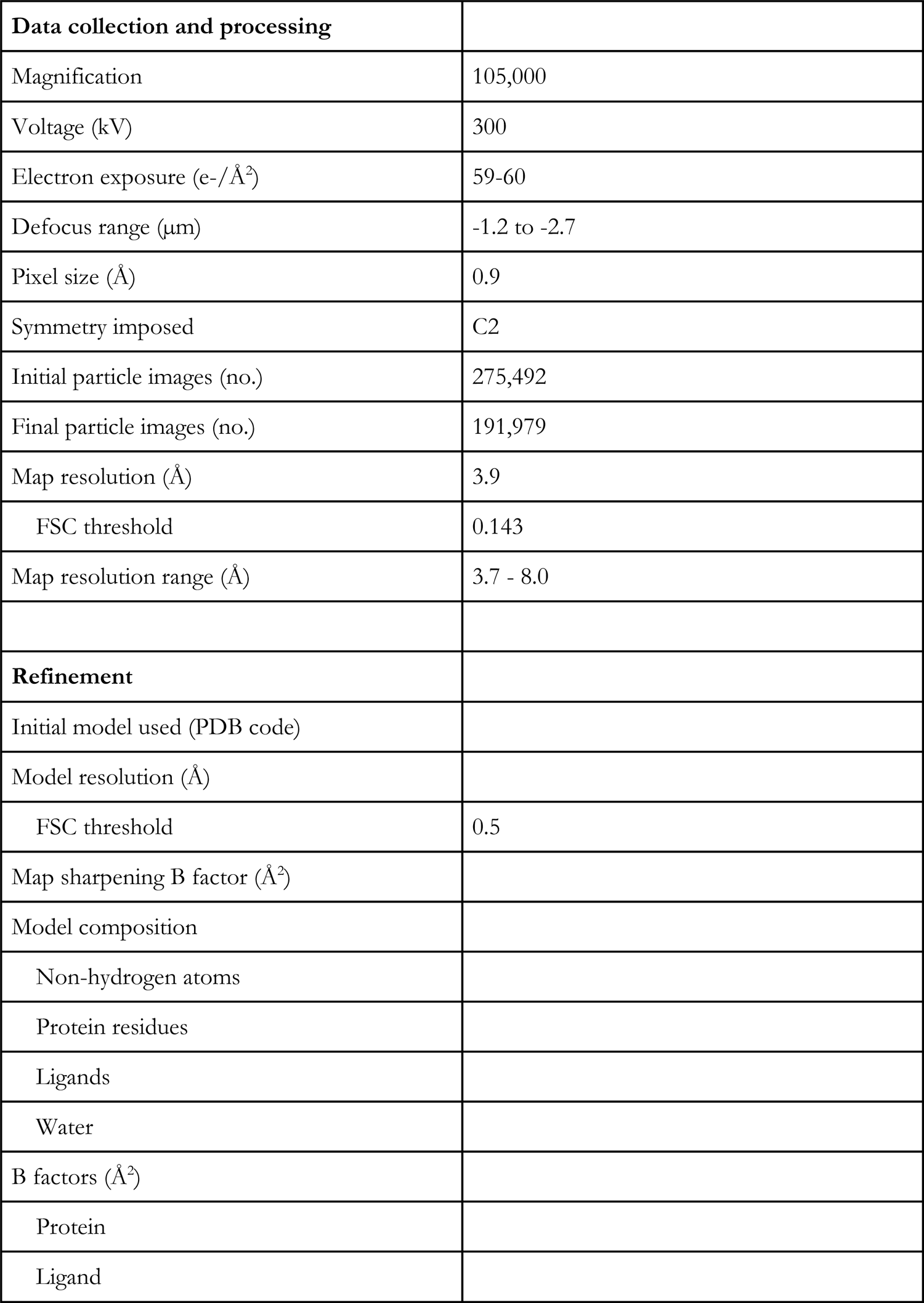
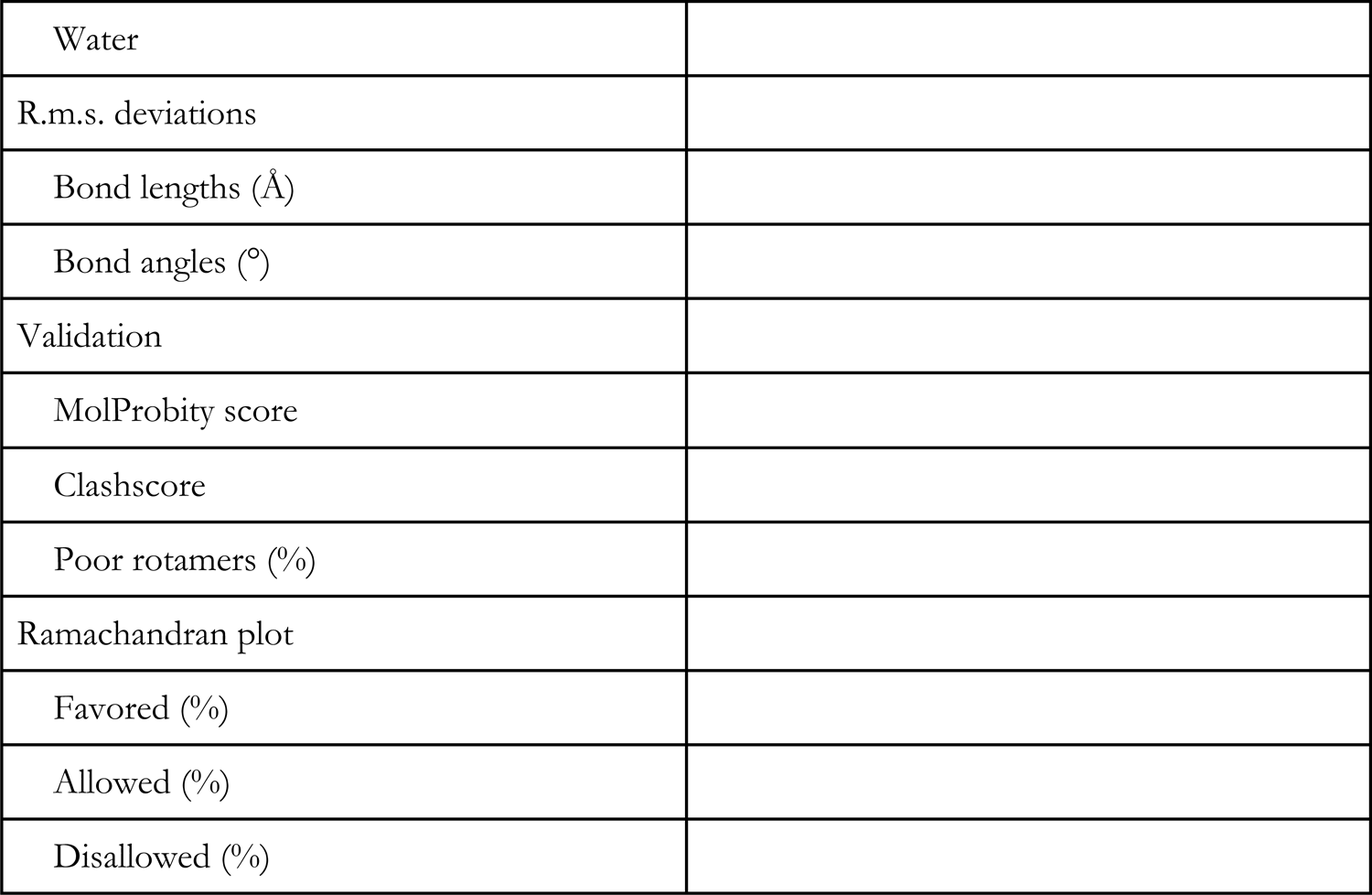
Cryo-EM data collection, refinement and validation statistics

|  |  |
| --- | --- |
| **Data collection and processing** |  |
| Magnification | 105,000 |
| Voltage (kV) | 300 |
| Electron exposure (e-/Å<sup>2</sup>) | 59-60 |
| Defocus range (μm) | -1.2 to -2.7 |
| Pixel size (Å) | 0.9 |
| Symmetry imposed | C2 |
| Initial particle images (no.) | 275,492 |
| Final particle images (no.) | 191,979 |
| Map resolution (Å) | 3.9 |
| FSC threshold | 0.143 |
| Map resolution range (Å) | 3.7 - 8.0 |
| **Refinement** |  |
| Initial model used (PDB code) |  |
| Model resolution (Å) |  |
| FSC threshold | 0.5 |
| Map sharpening B factor (Å<sup>2</sup>) |  |
| Model composition |  |
| Non-hydrogen atoms |  |
| Protein residues |  |
| Ligands |  |
| Water |  |
| B factors (Å<sup>2</sup>) |  |
| Protein |  |
| Ligand |  |

|  |
| --- |
| Water |
| R.m.s. deviations |
| Bond lengths (Å) |
| Bond angles (°) |
| Validation |
| MolProbity score |
| Clashscore |
| Poor rotamers (%) |
| Ramachandran plot |
| Favored (%) |
| Allowed (%) |
| Disallowed (%) |

## Supplemental Figure Legends

**Figure 1 – supplement 1.**
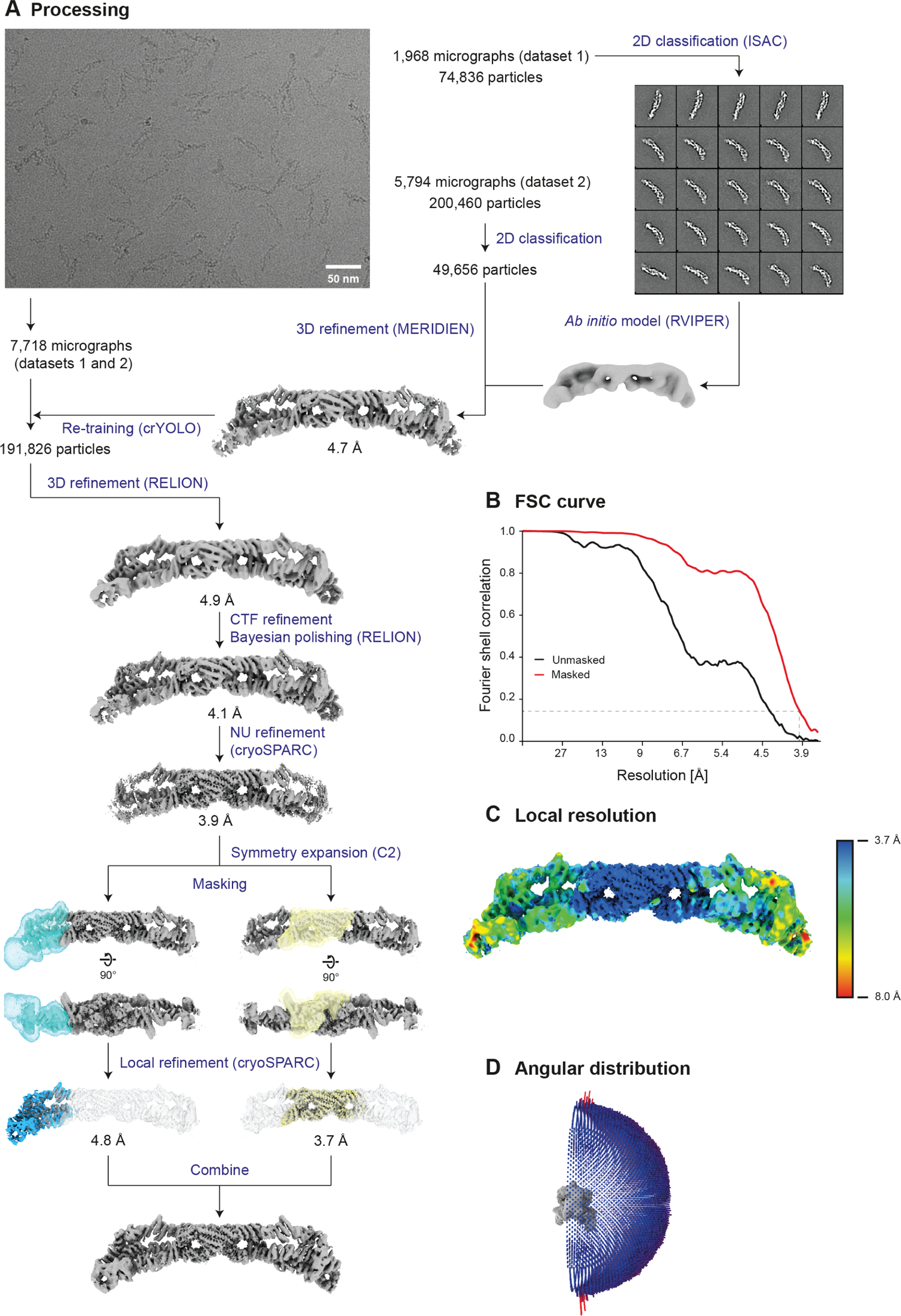
EM data analyses. (**A**) Processing scheme including an exemplary micrograph and a subset of selected 2D classes of RZZ. (**B**) Fourier Shell Correlation (FSC) curves of a global (i.e. non-focused) non-uniform 3D refinement in cryoSPARC. The dashed line indicates the 0.143 FSC criterion that intersects the masked FSC curve at 3.94 Å. (**C**) Local resolution plotted on the locally filtered reconstruction obtained by non-uniform refinement in a rainbow-colored gradient from blue (3.7 Å) to red (8.0 Å). (**D**) Angular distribution displayed as colored bars.

**Figure 1 – supplement 2.**
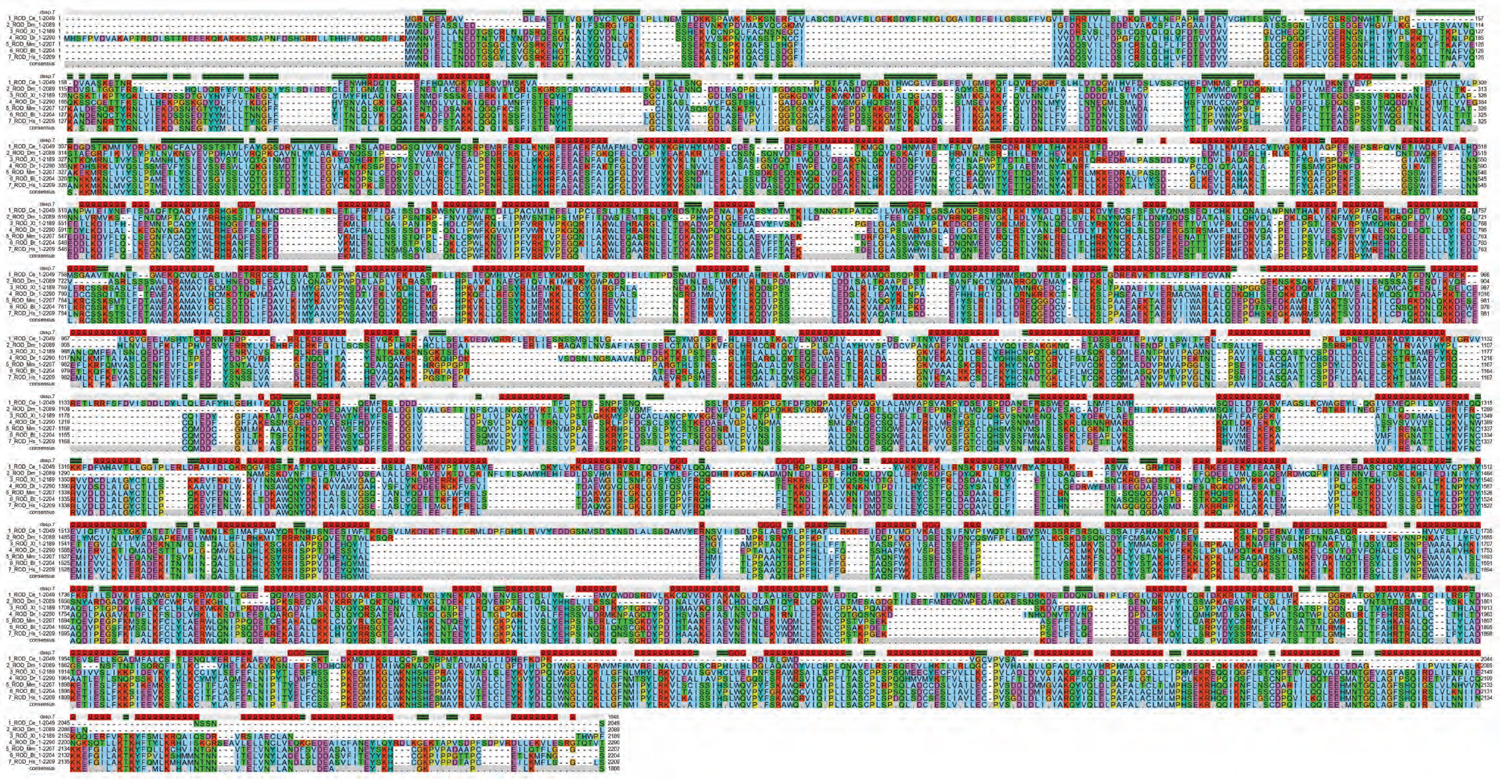
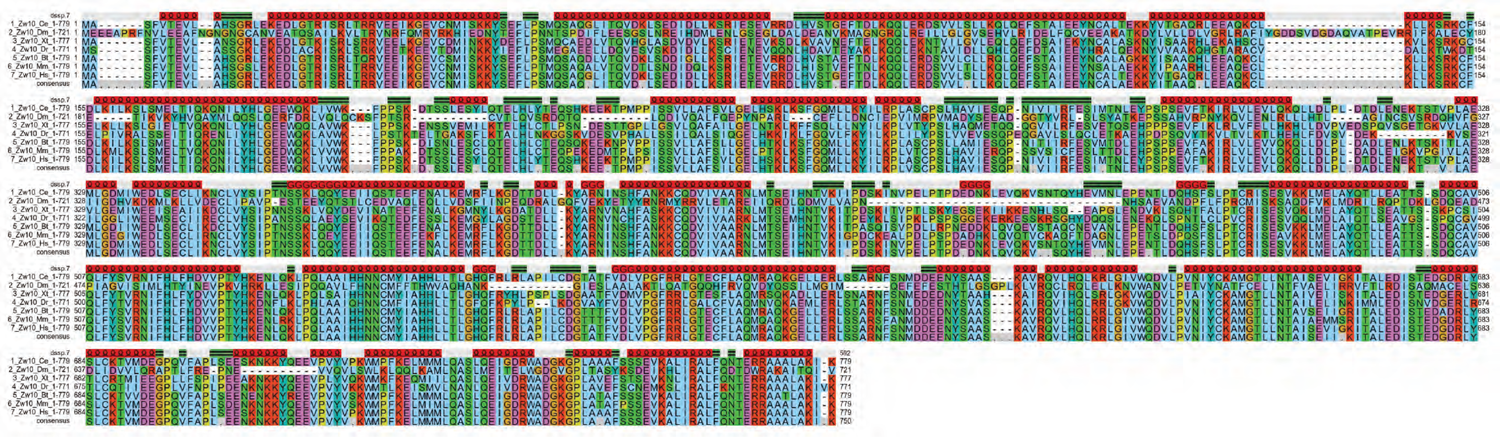
Multiple sequence alignment of ROD and ZW10 ROD and ZW10 sequences from *Caenorhabditis elegans* (Ce), *Drosophila melanogaster* (Dm), *Xenopus tropicalis* (Xt), *Danio rerio* (Dr), *Bos taurus* (Bt), *Mus musculus* (Mm), and *Homo sapiens* (Hs) were aligned with MAFFT (Katoh et al., 2002) and visualized with software developed in house. The secondary structure of the two RZZ subunits (straight black lines on green, *β*-strands; loopy black lines on red, helices; grey, coils) is displayed above the aligned sequences. For ROD, note in the second row a helical hairpin discussed in the text that lines the farnesyl-binding cavity, and the corresponding short deletions in species (Ce and Dm) where Spindly is not farnesylated.

**Figure 2 – Supplement 1.**
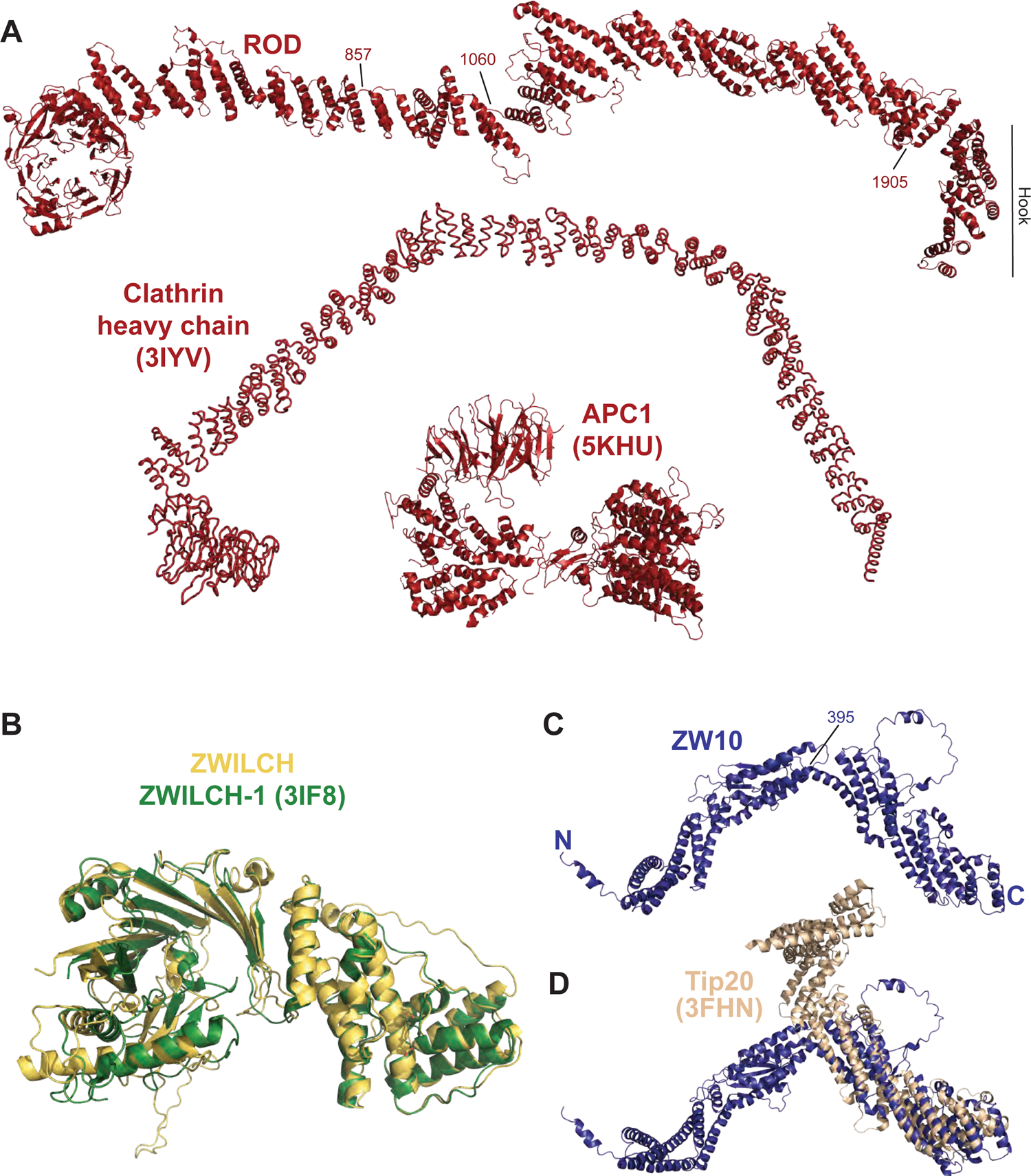
Comparison of structural homologs of RZZ subunits. (**A**) Cartoon model of ROD and of the indicated proteins that share the same structural organization of ROD. PDB ID codes are included. (**B**) Structural superposition of Zwilch in our cryo-EM reconstruction (yellow) and of the previously published crystal structure of Zwilch (green) (Civril et al., 2010). (**C**) Cartoon model of ZW10 and superposition with Tip20, a related S. cerevisiae’s ortholog. The interdomain angle of the N- and C-terminal domains is very different in the two structures.

**Figure 3 – Supplement 1.**
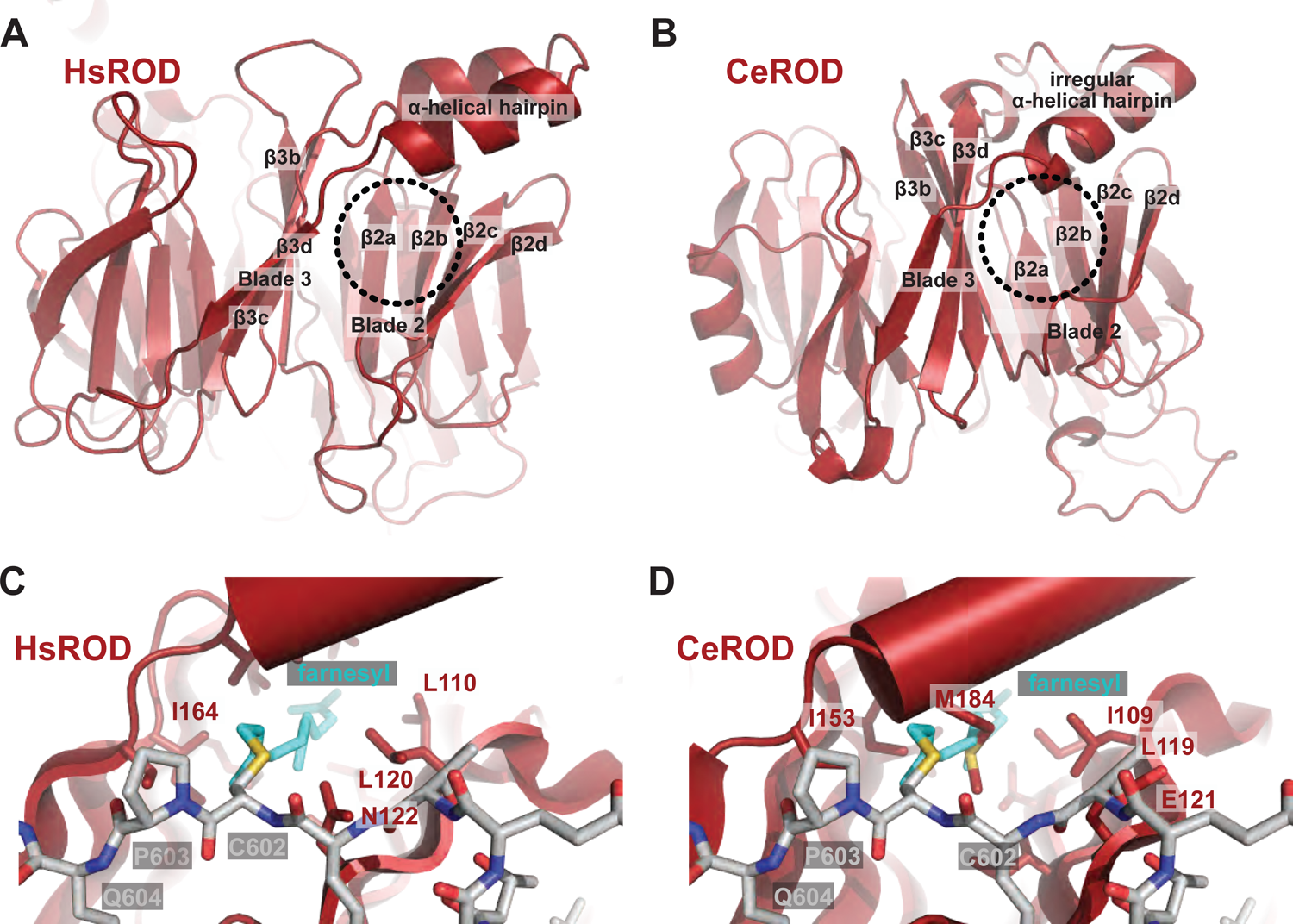
Comparison of the HsROD and CeROD β-propellers. (**A**) The *β*-propeller of human ROD already shown in Figure 3B with the indicated farnesyl-binding site. (**B**) *β*-propeller of *C. elegans*’ ROD predicted by AlphaFold2 (Jumper et al., 2021; Tunyasuvunakool et al., 2021). (**C**-**D**) Modelled farnesyl groups in the two structures demonstrate occlusion of the binding cavity in *C. elegans*, where M184 is predicted to sterically clash with a hypothetical farnesyl moiety.

**Figure 4 – Supplement 1.**
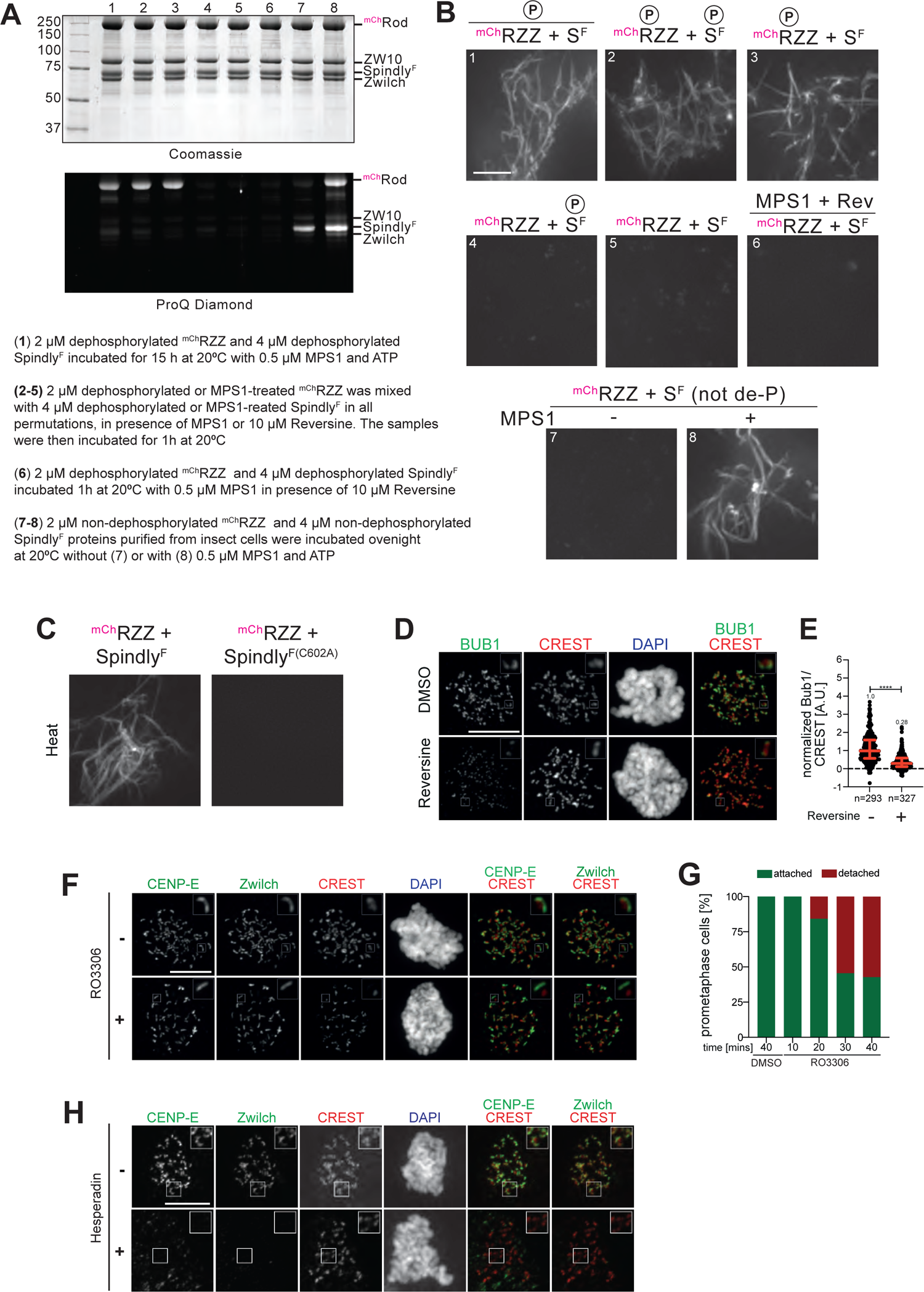
Additional polymerization and cell biology experiments. (**A-B**) Assessment of requirements for MPS1 phosphorylation for corona assembly *in vitro*. Samples 1-8 are shown on the left in Coomassie-stained and ProQ Diamond-stained SDS-PAGE. The content of samples 1-8 is described in the legend under panel A, and additionally for samples 2-5 over each polymerization experiment in panel B, where the encircled P signals which sample was pre-phosphorylated with MPS1 and which samples were treated with reversine to inhibit MPS1. In B, samples were imaged in a spinning disk confocal microscope at 561 nm. Scale bar = 5 µm. (**C**) Heat-induced polymerization experiments with ^mCh^RZZ and wild-type Spindly^F^ or Spindly^F(C602A)^ that cannot be farnesylated. The same positive control is also shown in Figure 7 – Supplement 1B. (**D**) Levels of BUB1 at kinetochores of HeLa cells that had been previously synchronized in G2 phase with 9 μM RO3306 for 16 h and then released into mitosis. Subsequently, cells were immediately treated with 500 nM reversine, 3.3 µM nocodazole, and 10 µM MG132 for 1 hour. CREST serum was used to visualize kinetochores and DAPI to stain DNA. Scale bar: 10 µm. (**E**) Scatter dot plots representing normalized intensity ratios of BUB1 over CREST for individual kinetochores of cells from the experiment shown in panel C. Red lines indicate median with interquartile range. (**F**) HeLa cells were synchronized in G2 phase with 9 µM RO3306 for 16 hours and the released into mitosis by withdrawing the inhibitor. Cells were immediately treated with 3.3 µM nocodazole to prevent microtubule depolymerization and allow maximal corona expansion. After one hour, cells were treated again with RO3306 for the indicated time points, before fixation and further processed for fluorescence microscopy. CREST serum was used to visualize kinetochores and DAPI to visualize DNA. Scale bar: 10 µm. (**G**) Quantification of the experiment in panel E. (**H**) DLD-1 cells treated like in panel E were released from the G2 arrest into mitosis for 1 hour in presence of 3.3 µM nocodazole, 500 nM hesperadine, and 10 µM MG132, fixed, and further processed for fluorescence microscopy.

**Figure 5 – Supplement 1.**
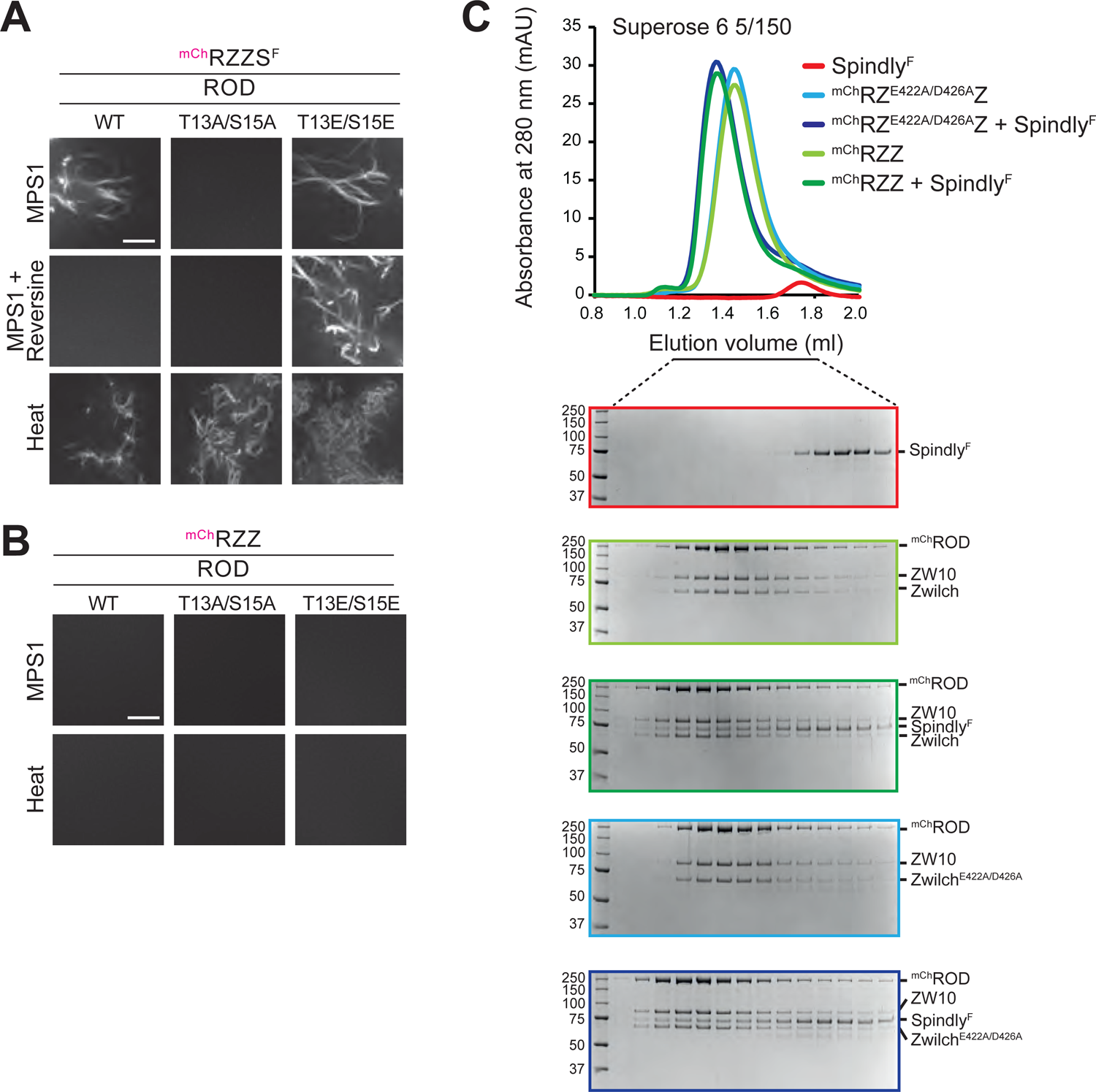
Additional in vitro polymerization and biochemical experiments. (**A**) The first two rows of experiments are already displayed as the two rows of Figure 5A. The third row is added to demonstrate that the ^mCh^RZZS^F^ complex carrying T13A/S15A mutations on ROD, which does not spontaneously filament in presence of MPS1 at 20°C, will form filaments upon mild heating at 30°C, indicating that its ability to polymerize is not compromised. Scale bar: 5 µM. (**B**) None of the indicated samples forms filaments in absence of Spindly^F^. Scale bar: 5 µm. (**C**) Size-exclusion chromatography experiment demonstrating that RZZ complex carrying E422A/D426A mutations interacts with Spindly^F^ as strongly as its wild type counterpart.

**Figure 6 – Supplement 1.**
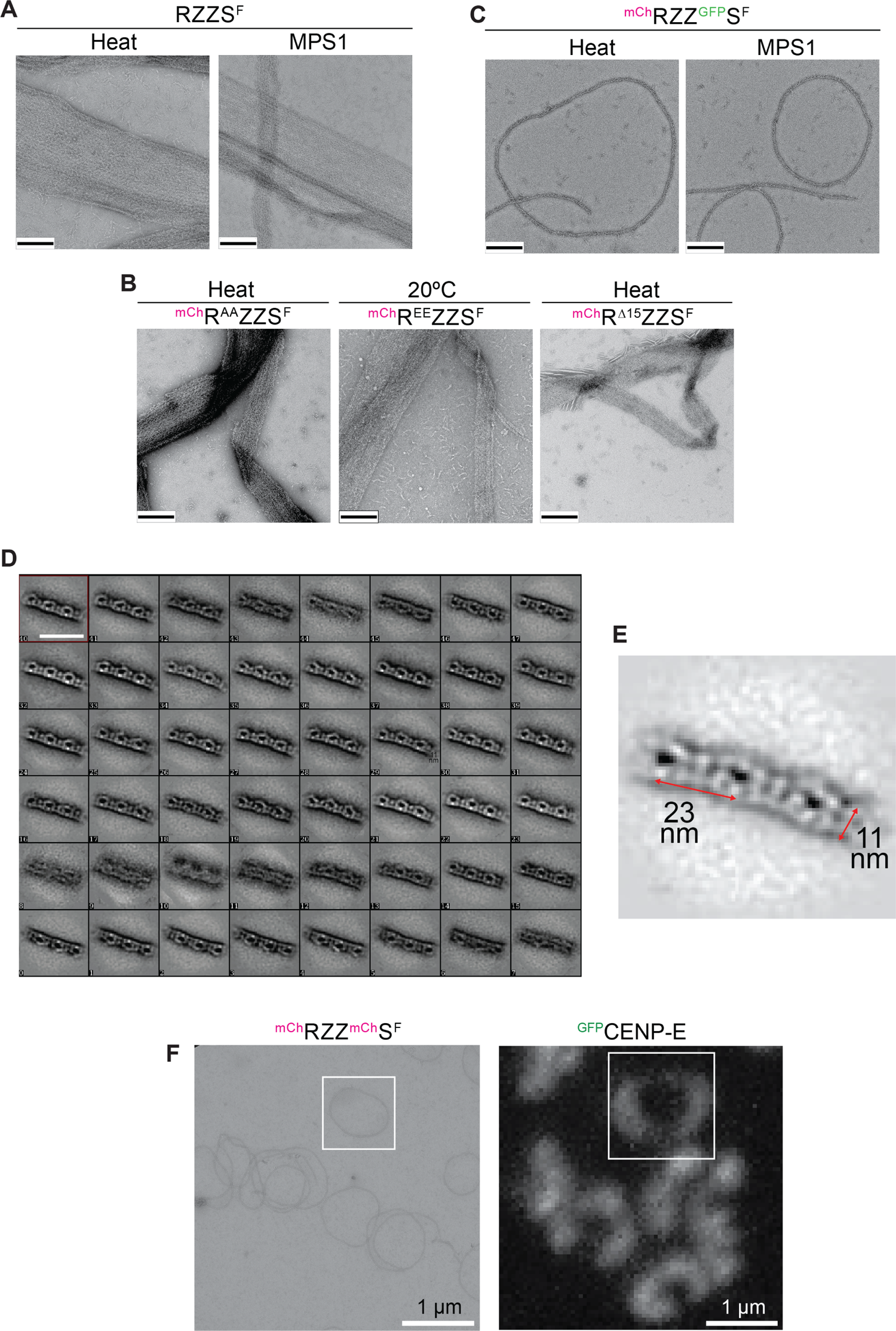
(**A**) Untagged RZZS^F^ formed sheets indistinguishable from those formed by ^mCh^RZZS^F^. Scale bar (black line): 200 nm. (**B**) Sheets were also formed by ^mCh^RZZS^F^ containing the T13A/S15A mutations on ROD upon heating the sample to 30°C. The phosphomimetic mutant T13E/S15E of the same complex assembled into filaments spontaneously at 20°C. The ^mCh-Δ15^RZZS^F^ forms sheets at 30°C. Scale bar (black line): 200 nm. (**C**) ^mCh^RZZ/^GFP^S^F^ formed ring structures or curved filaments similar to those observed with ^mCh^RZZ/^mCh^S^F^. (**D**) 2D class averages of segments of negatively-stained single circles or filaments like those shown in panel C. Note the extreme orientation preference of the various classes. Scale bar: 50 nm. (**E**) Enlargement of one class average shown in D with indicated dimensions. (**F**) A size comparison from a field of ^mCh^RZZ^mCh^S^F^ and ^GFP^CENP-E at kinetochore coronas of nocodazole-treated HeLa cells is shown ad the same magnification to emphasize the similarity of curvatures in rings and coronas.

**Figure 7 – Supplement 1.**
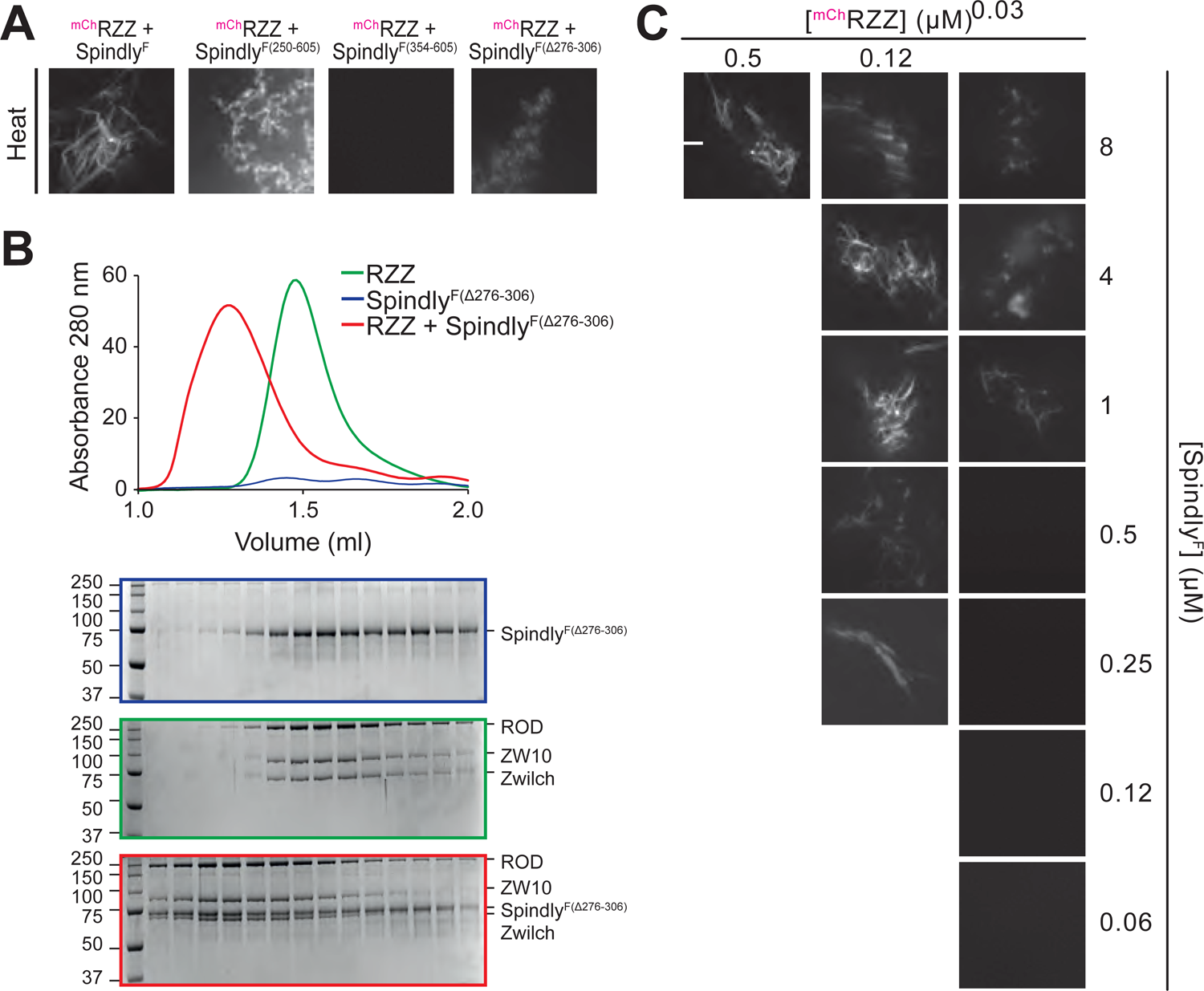
Additional biochemical experiments. (**A**) Heat-induced filaments of the indicated species. Note that Spindly^F(250-605)^ and Spindly^F(Δ276-306)^ may show a combination of slight precipitation and filamentation when filamentation is induced with heat. The same positive control is also shown in Figure 5 – Supplement 1D. (**B**) Size-exclusion chromatography of Spindly^F(Δ276-306)^, RZZ, and their complex. (**C**) Titration of ^mCh^RZZ and Spindly^F^ at the indicated concentrations.

